# Single cell chromatin accessibility reveals pancreatic islet cell type- and state-specific regulatory programs of diabetes risk

**DOI:** 10.1101/693671

**Authors:** Joshua Chiou, Chun Zeng, Zhang Cheng, Jee Yun Han, Michael Schlichting, Serina Huang, Jinzhao Wang, Yinghui Sui, Allison Deogaygay, Mei-Lin Okino, Yunjiang Qiu, Ying Sun, Parul Kudtarkar, Rongxin Fang, Sebastian Preissl, Maike Sander, David Gorkin, Kyle J Gaulton

## Abstract

Genetic risk variants for complex, multifactorial diseases are enriched in *cis*-regulatory elements. Single cell epigenomic technologies create new opportunities to dissect cell type-specific mechanisms of risk variants, yet this approach has not been widely applied to disease-relevant tissues. Given the central role of pancreatic islets in type 2 diabetes (T2D) pathophysiology, we generated accessible chromatin profiles from 14.2k islet cells and identified 13 cell clusters including multiple alpha, beta and delta cell clusters which represented hormone-producing and signal-responsive cell states. We cataloged 244,236 islet cell type accessible chromatin sites and identified transcription factors (TFs) underlying both lineage- and state-specific regulation. We measured the enrichment of T2D and glycemic trait GWAS for the accessible chromatin profiles of single cells, which revealed heterogeneity in the effects of beta cell states and TFs on fasting glucose and T2D risk. We further used machine learning to predict the cell type-specific regulatory function of genetic variants, and single cell co-accessibility to link distal sites to putative cell type-specific target genes. We localized 239 fine-mapped T2D risk signals to islet accessible chromatin, and further prioritized variants at these signals with predicted regulatory function and co-accessibility with target genes. At the *KCNQ1* locus, the causal T2D variant rs231361 had predicted effects on an enhancer with beta cell-specific, long-range co-accessibility to the insulin promoter, and deletion of this enhancer reduced insulin gene and protein expression in human embryonic stem cell-derived beta cells. Our findings provide a cell type- and state-resolved map of gene regulation in human islets, illuminate likely mechanisms of T2D risk at hundreds of loci, and demonstrate the power of single cell epigenomics for interpreting complex disease genetics.

## Introduction

Gene regulatory programs are largely orchestrated by *cis*-regulatory elements that direct the expression of genes in response to specific developmental and environmental cues. Genetic variants associated with disease by genome-wide association studies (GWAS) are highly enriched within putative *cis-*regulatory elements^1^, highlighting the importance of regulatory sequence in mediating disease risk. The activity of regulatory elements is often restricted to specific cell types and/or cell states, limiting the ability of ATAC-seq and other “ensemble” (or “bulk”) epigenomic technologies to map regulatory elements in individual cell types within disease-relevant tissues. To overcome this limitation, new approaches to obtain ATAC-seq profiles from single nuclei (snATAC-seq) allow for the disaggregation of open chromatin from heterogenous samples into component cell types and subtypes^2–5^. These developments create opportunities to dissect the molecular mechanisms that underlie genetic risk of disease. However, to date snATAC-seq data from disease-relevant human tissues are limited^6–9^.

Type 2 diabetes (T2D) is a multifactorial disease with a highly polygenic inheritance^10^. Pancreatic islets are central to genetic risk of T2D, as evidenced by shared association between T2D risk and quantitative measures of islet function^11–13^ and enrichment of T2D risk variants in islet regulatory sites^14–18^. Islets are comprised of multiple endocrine cell types with distinct functions^19–21^ and are heterogeneous^22–24^ in gene expression and other molecular signatures which likely reflect different functional cell states^22,25,26^. Heterogeneity in the epigenome of islet cell types has not been described, however, which is necessary to understand islet regulation and interpret the molecular mechanisms of non-coding T2D risk variants. In this study, we map accessible chromatin profiles of individual islet cells using snATAC-seq, define the regulatory programs of islet cell types and cell states, describe their relationship to T2D risk and fasting glycemia, and predict the molecular mechanisms of T2D risk variants.

## Results

### Islet snATAC-seq reveals 13 cell clusters with distinct regulatory landscapes

To map the accessible chromatin landscape of single islet cells, we performed snATAC-seq on human pancreatic islets from three donors (Supplementary Table 1). We used a combinatorial barcoding snATAC-seq approach previously optimized by our group for use on tissues^2,4^ (see Methods). To confirm library quality, we first analyzed the data as ensemble ATAC-seq by aggregating all high-quality mapped reads irrespective of barcode. Ensemble snATAC-seq from all three samples showed the expected insert size distribution (Supplementary Figure 1a), strong enrichment of signal at transcription start sites (TSS) (Supplementary Figure 1b), and high concordance of signal with published islet ATAC-seq data^14,27–29^ (Supplementary Figure 1c).

To obtain a collection of high-quality single cell profiles, we first filtered out cells with less than 1,000 reads (Supplementary Figure 1d), resulting in a total of 17,995 cells across the three samples. We then clustered accessible chromatin profiles from these cells, making key modifications to previous approaches (see Methods for details)^4^. First, as the inputs to clustering we used normalized read counts in 5 kb sliding windows genome-wide rather than read counts within ensemble peak calls, reasoning that ensemble peak calls could be biased towards more common cell types. Second, we performed an initial round of clustering and quality control on a per-sample basis, which removed 2,709 cells in low read depth clusters. Third, prior to clustering cells across samples, we used mutual nearest neighbors^30^ to correct for variability across donors. Finally, we clustered all cells together and performed additional quality control by removing one cluster without representation from all donors (694 cells), and one with aberrant read depth and low intra-cluster similarity (192 cells). After all clustering and filtering steps, we retained 14,239 cells which mapped to 13 clusters, all of which had consistent representation across samples and read depth profiles (Figure 1a, Supplementary Figure 2a-c).

**Figure 1.**
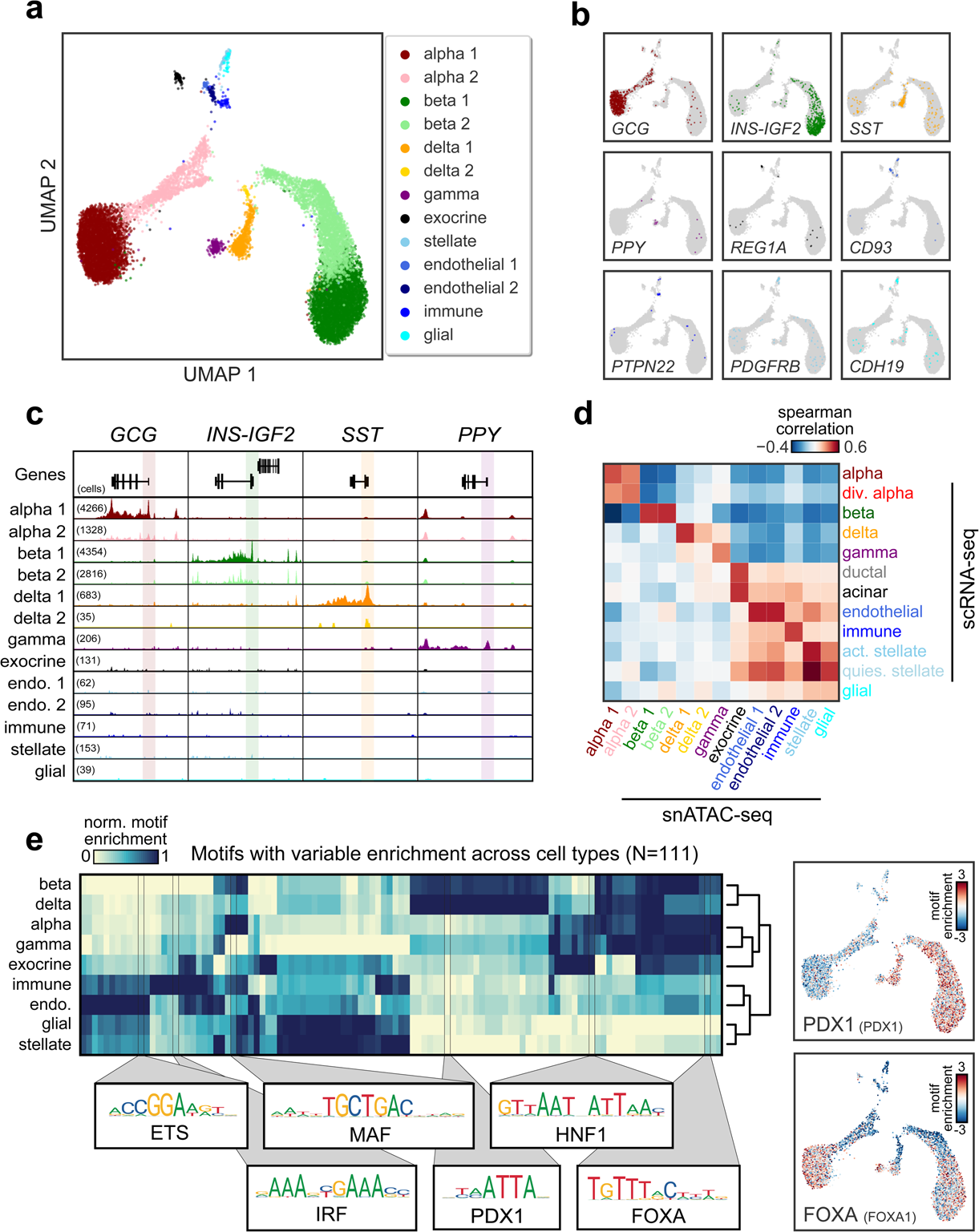
Pancreatic islet cell type accessible chromatin defined using snATAC-seq. (a) Clustering of accessible chromatin profiles from 14,239 pancreatic islet cells identifies 13 distinct clusters. Cells are plotted using the first two UMAP components, and clusters are assigned cell type identities based on promoter accessibility of known marker genes for each cell type. (b) Promoter accessibility in a 1 kb window around the TSS for selected endocrine and non-endocrine marker genes for each profiled cell. A cell is colored if it had promoter accessibility for the marker gene listed in the bottom right corner of each subplot, and otherwise is grey. (c) Genome browser plots showing aggregate read density (scaled to uniform 1×10^5^ read depth, range: 1-10) for cells within each cell type cluster at hormone gene loci for endocrine islet cell types: *GCG* (alpha), *INS-IGF2* (beta), *SST* (delta), and *PPY* (gamma). The promoter region for each gene is highlighted, and the number of cells for each cell type cluster is listed in parenthesis. (d) Spearman correlation between t-statistics of marker genes based on promoter accessibility (snATAC-seq) or gene expression (scRNA-seq) using the top 100 most specific gene promoters from each islet snATAC-seq cluster. (e) Normalized chromVAR motif enrichment values for 111 TF sequence motifs that have variable activity across clusters. We collapsed multiple clusters for each cell type into a single cluster (e.g. combining beta 1 and beta 2 into a single beta cell cluster). Subtype-specific motif enrichment is presented in Figure 2. Position weight matrices and names are shown for sequence motifs for TF families enriched across different endocrine and non-endocrine cell types. Enrichment z-scores for FOXA1 and PDX1 motifs in each cell are projected onto UMAP coordinates to the right of the main heatmap.

To determine the cell type represented by each cluster, we examined chromatin accessibility at the promoter region of the cognate hormone genes for endocrine cells and known marker genes for non-endocrine cell types. Based on these marker gene promoters, we identified clusters representing beta (*INS-IGF2/*insulin), alpha (*GCG/*glucagon), delta (*SST/*somatostatin), gamma (*PPY*/pancreatic polypeptide) cells, exocrine acinar and ductal (labeled as ‘exocrine’; *REG1A, S100A14*)^31,32^, immune (*PTPN22*)^32^, stellate (*PDGFRB*)^32^, glial (*CDH19*)^33^, and endothelial (*CD93*)^34^ cells (Figure 1b-c, Supplementary Figure 2d). We defined a broader set of marker gene promoters for each cluster by identifying gene promoters with differential accessibility across clusters and retaining the top 100 differential promoters for each cluster (see Methods, Supplementary Table 2). To validate the cell type we assigned to each cluster, we derived gene expression marker genes from published islet scRNA-seq data^23^ and correlated t-statistics of snATAC-seq marker gene promoters with t-statistics of scRNA-seq marker genes (see Methods, Supplementary Figure 3a-e). We observed highly specific correlations between marker genes of endocrine and other pancreatic cell types in snATAC-seq and scRNA-seq (Figure 1d). Of note, the multiple clusters of alpha, beta, and delta cells in snATAC-seq each had strongest correlation with their respective cell type.

To characterize the regulatory programs of each cell type, we aggregated reads for cells within each cluster and identified accessible chromatin sites for the cluster using MACS2 (see Methods). In total we identified 244,236 accessible chromatin sites merged across the 13 clusters (Supplementary Data 1), which were concordant with sites identified in ensemble islets (Supplementary Figure 4a-b). Notably, accessible chromatin in alpha and beta cells was highly concordant with bulk ATAC-seq of corresponding FACS-sorted populations^35,36^, confirming that we identified cell type-specific islet chromatin (Supplementary Figure 4c). To next understand the regulatory logic underlying islet cell types, we used chromVAR^37^ to identify TF sequence motifs from JASPAR^38^ enriched within accessible chromatin of each cell. We focused on 111 TF motifs with evidence for variability across cells (see Methods, Supplementary Figure 4d, Supplementary Table 3). Analysis of motif enrichments averaged across cells for each cell type revealed distinct patterns of motif enrichment across cell types, many consistent with known functions in islet cells (Figure 1e, Supplementary Table 3). For example, the PDX1 motif was enriched in beta (normalized enrichment=0.93) and delta (1.0) cells^39^, and MAF motifs were enriched in alpha (1.0) and beta cells (0.93)^40–42^ (Figure 1e). We also identified motif enrichments shared across all endocrine cell types, such as FOXA, and in non-endocrine cell types, including IRF for immune^43^ (1.0) and ETS for endothelial^44^ (1.0) cells (Figure 1e). Hierarchical clustering of cell types based on TF motif enrichment patterns further revealed that regulatory programs of beta and delta cells were closely related as were the programs of alpha and gamma cells (Figure 1e), consistent with single cell expression data^31,32,45^.

### Heterogeneity in islet endocrine cell accessible chromatin and regulatory programs

A major strength of single cell approaches is the ability to reveal heterogeneity within a cell type. Indeed, our initial clustering showed that alpha, beta and delta cells segregated into sub-clusters. To characterize these sub-clusters, we determined gene promoter accessibility in each sub-cluster and identified promoters with variable accessibility between sub-clusters (see Methods, Supplementary Data 2). We focused on alpha and beta cells, where cell numbers allowed for robust calculations. Notably, we found *INS* among genes with the most variable promoter accessibility between beta cell sub-clusters (*INS-IGF2* beta OR=5.05, two-sided Fisher’s exact P=3.98×10^−37^), leading us to rename the beta 1 and beta 2 clusters as INS^high^ and INS^low^ beta cells, respectively (Figure 1b-c; Figure 2a). Similarly, *GCG* promoter accessibility was highly variable between alpha cell sub-clusters (*GCG* alpha OR=3.30, P=4.68×10^−25^), and we renamed the alpha 1 and alpha 2 sub-clusters as GCG^high^ and GCG^low^ alpha cells, respectively (Figure 1b-c; Figure 2a).

**Figure 2.**
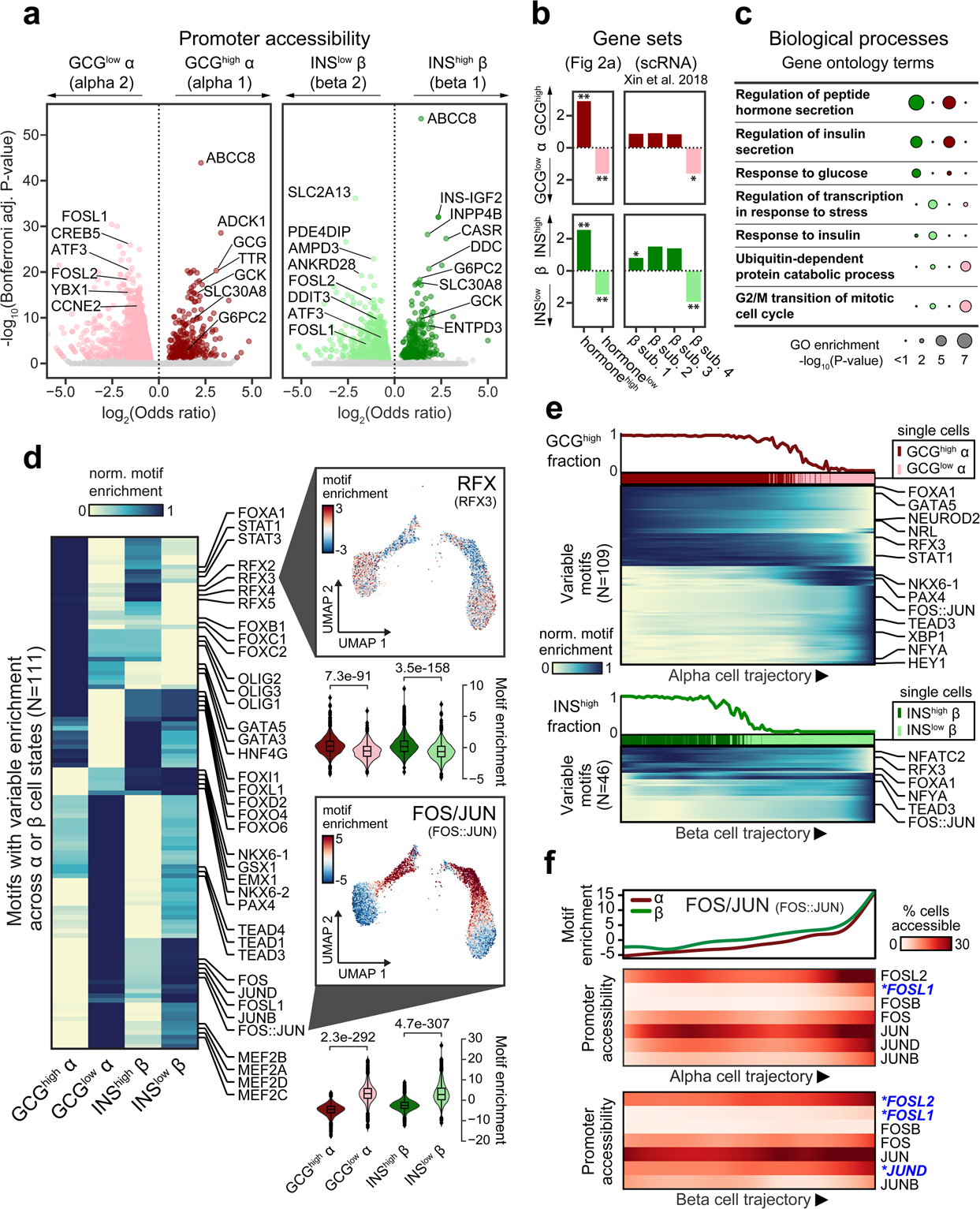
Heterogeneity in alpha and beta cell accessible chromatin and regulatory programs. (a) Gene promoters with significantly differential chromatin accessibility between sub-clusters of alpha cells (left) and beta cells (right). Genes with increased promoter accessibility in alpha 1 (GCG^high^) and beta 1 (INS^high^) sub-clusters include *GCG* (glucagon) for alpha cells and *INS* (insulin) for beta cells, as well as genes such as *ABCC8, G6PC2, GCK* and *SLC30A8.* Conversely, genes with increased promoter accessibility in alpha 2 (GCG^low^) and beta 2 (INS^low^) sub-clusters include genes such as *FOSL1, FOSL2*, and *ATF3.* (b) Genes with increased promoter accessibility in the hormone-high (INS^high^, GCG^high^) or hormone-low (INS^low^, GCG^low^) state of one cell type were significantly enriched for genes with increased hormone-high or hormone-low activity in the other cell type, respectively (left); Genes with differential promoter accessibility across alpha and beta cell states were enriched for genes in beta cell subsets (β sub. 1-4) previously identified in an islet single cell gene expression study. (right) **FDR<.01, *FDR<.10. (c) Gene ontology terms for biological processes related to glucose response and hormone secretion were enriched in genes with higher promoter accessibility in INS^high^ and GCG^high^ cells, whereas terms for stress response, insulin signaling and cell cycle were enriched in genes with higher promoter accessibility in INS^low^ and GCG^low^ cells. (d) Row-normalized chromVAR enrichments for 111 TF motifs showing variable enrichment across alpha or beta cells. We observed motifs enriched for different sub-clusters including RFX family members (RFX2-5) for GCG^high^ alpha and INS^high^ beta cells, and FOS/JUN family members for GCG^low^ alpha and INS^low^ beta cells. Individual cell enrichment z-scores of a representative RFX (RFX3) and FOS/JUN (FOS::JUN) motif are plotted on UMAP coordinates, and the violin plots below each UMAP plot show enrichment values (median: center line, boxplot limits: quartiles) within each alpha and beta state. (e) Ordering of alpha and beta cells on a trajectory using high *GCG/INS-IGF2* promoter accessibility as the anchor point with Cicero. Plots show cells binned across this trajectory from left to right, where the top shows the percentage of cells in the hormone-high state in a given bin, colored bars above the heatmap represent individual cells with their binary clusters in their positions across each trajectory, and the heatmap shows chromVAR enrichments for motifs in bins across each trajectory. (f) Motifs in the FOS/JUN family show increasing enrichment across the alpha and beta cell trajectory. Genes in the FOS/JUN family with promoter accessibility patterns that match the motif enrichment patterns (Spearman correlation>.9) are highlighted (in blue and starred).

Apart from *INS* and *GCG*, we found significant overlap in the genes that distinguish hormone-high (INS^high^, GCG^high^) from hormone-low (INS^low^, GCG^low^) alpha and beta cells by gene set enrichment analysis (GSEA) (Figure 2b). Genes with increased promoter accessibility in hormone-high states including *GCK, ABCC8, G6PC2* and *SLC30A8* were enriched for processes such as hormone secretion and glucose response (Figure 2a,c, Supplementary Table 4). In contrast, genes with increased promoter accessibility for hormone-low states including *ATF3*, *FOSL1*, and *FOSL2* and were enriched for stress-induced signaling response^46^ (Figure 2a,c, Supplementary Table 4). Similar states were also evident in delta cells, although low cell numbers impede deeper analysis in our study (Supplementary Figure 5). We compared genes with significantly different promoter accessibility between states to gene sets describing beta cell heterogeneity (β-sub.1-4) from a previous scRNA-seq study^23^. Genes with increased promoter accessibility in hormone-low cells (INS^low^, GCG^low^) were enriched in a beta cell sub-cluster (β-sub.4) associated with ER stress and protein folding and with low *INS* expression, whereas genes with increased promoter accessibility in hormone-high cells (INS^high^, GCG^high^) were enriched in the other beta cell sub-clusters (β-sub.1-3) (Figure 2b). These data reveal epigenomic differences between endocrine cell states among genes involved in hormone production and stress-induced signaling responses, and point to an underlying commonality in the genes that govern state-specific functions across different endocrine cell types.

The transcriptional regulatory programs driving functional heterogeneity in alpha and beta cells are unknown. Therefore, we determined TF sequence motifs differentially enriched across alpha and beta cell states. We focused on 111 TF motifs showing evidence for variable enrichment between alpha and beta cell states (see Methods, Supplementary Figure 6a, Supplementary Table 5) and observed clear patterns that distinguished different states within alpha and beta cells, again revealing commonalities across cell types (Figure 2d). For example, motifs for RFX family members were enriched in hormone-high states (GCG^high^, INS^high^), but not in hormone-low states (GCG^low^, INS^low^) (RFX3 - mean INS^high^ enrich=.26, INS^low^=-.62, P=3.5×10^−158^; GCG^high^=.29, GCG^low^=-.56, P=7.3×10^−91^) (Figure 2d). In contrast, motifs for FOS and JUN family members were prominently enriched in hormone-low states, but not the hormone-high states (FOS::JUN - mean INS^high^ enrich=-1.45, INS^low^=4.50, P=4.7×10^−307^; GCG^high^=-1.45, GCG^low^=4.46, P=2.3×10^−292^) (Figure 2d). Again, we also observed similar motif enrichment patterns between delta cell states (Supplementary Figure 6a-b).

Analysis of single cells ordered along a trajectory has been used to examine gene regulatory programs as a continuum rather than as discrete or binary states^6,23,47^. To explore potential gradations among alpha and beta cells, we used Cicero^6^ to order alpha and beta cells along trajectories based on chromatin accessibility. We ordered cells using high promoter accessibility at *INS* (beta) or *GCG* (alpha) as the root states for each trajectory (see Methods). We refer to the axis of these trajectories as “pseudo-state” rather than the conventional “pseudo-time”, because the heterogeneity appears to be more related to cell state than to time. We observed cells on a gradient between hormone-high and hormone-low states of alpha and beta cells, and we noted a discernable transition point within the trajectory (Figure 2e, Supplementary Figure 7a-b). These trajectories allowed us to examine gene promoter accessibility and TF motif enrichment as a function of pseudo-state (Figure 2e, Supplementary Figure 7c). Consistent with the above results, lineage-specifying genes and enrichments for motifs in TF families such as RFX, Neurogenin-ATO and NFAT decreased along the trajectory from hormone-high to -low cells, whereas enrichment for motifs in TF families such as FOS/JUN, XBP and CCAAT (NFYA) increased along the trajectory (Figure 2e).

Structurally-related TFs often have similar motifs, and thus to assign motifs to specific TFs we correlated promoter-accessibility of TFs within the structural subfamily with motif enrichments across the state trajectory (see Methods)^48^. Motif enrichment for the FOS/JUN family correlated with the promoter accessibility of *FOSL1, FOSL2* and *JUND* across cells (Figure 2f), supporting a role for these specific TFs in hormone-low cell regulation. Similarly, motif enrichment for the Neurogenin-ATO subfamily correlated with promoter accessibility of *NEUROD1,* supporting a role for this TF in hormone-high cell regulation (Supplementary Figure 8a). While we did not observe strong correlations between RFX motif enrichment and promoter accessibility of *RFX* genes, the overall high promoter accessibility of *RFX6* and *RFX3* and known function in endocrine cells^49–51^ suggests they are TFs likely involved in hormone-high cell regulation (Supplementary Figure 8b).

### Enrichment of islet cell type- and state-specific regulatory sequences for diabetes- and fasting glycemia-associated genetic variants

Variants associated with complex diseases and physiological traits are enriched within *cis*- regulatory sequences^1,52^. More specifically, genetic variants influencing diabetes and fasting glucose level are enriched in pancreatic islet regulatory elements^15–17,53^. However, these enrichments based on ensemble data obscure the potential role of islet cell type- and state-specific regulation in these traits. Using our islet cell type- and state-resolved accessible chromatin profiles, we sought to determine the enrichment of genetic variants associated with type 1 and 2 diabetes^10,54^ and diabetes-related quantitative phenotypes^13,55–59^ as well as other complex traits and disease for calibration^60–67^. We first determined the enrichment of variants in accessible chromatin sites for each islet cell type and state using stratified LD score regression^68,69^ (see Methods). We observed significant enrichment (FDR<.1) of fasting glucose (FG) level and T2D association for both INS^high^ and INS^low^ beta cell states (T2D INS^high^ Z=4.45 q-value=.001, INS^low^ Z=4.00 q=.004; FG INS^high^ Z=3.93 q=.004, INS^low^ Z=3.34 q=.027), as well as enrichment of body-mass index (BMI) for SST^high^ delta cells (Z=3.50 q=.027) (Figure 3a). We also observed suggestive enrichment (P<.01) of 2hr glucose level adjusted for BMI for both alpha cell states (GCG^high^ Z=2.45 P=.007, GCG^low^ Z=2.40 P=.008), and T2D and fasting proinsulin level for GCG^low^ alpha cells (PI: Z=2.64, P=.004; T2D: Z=2.40 P=.008), although these enrichments did not pass multiple test correction.

**Figure 3.**
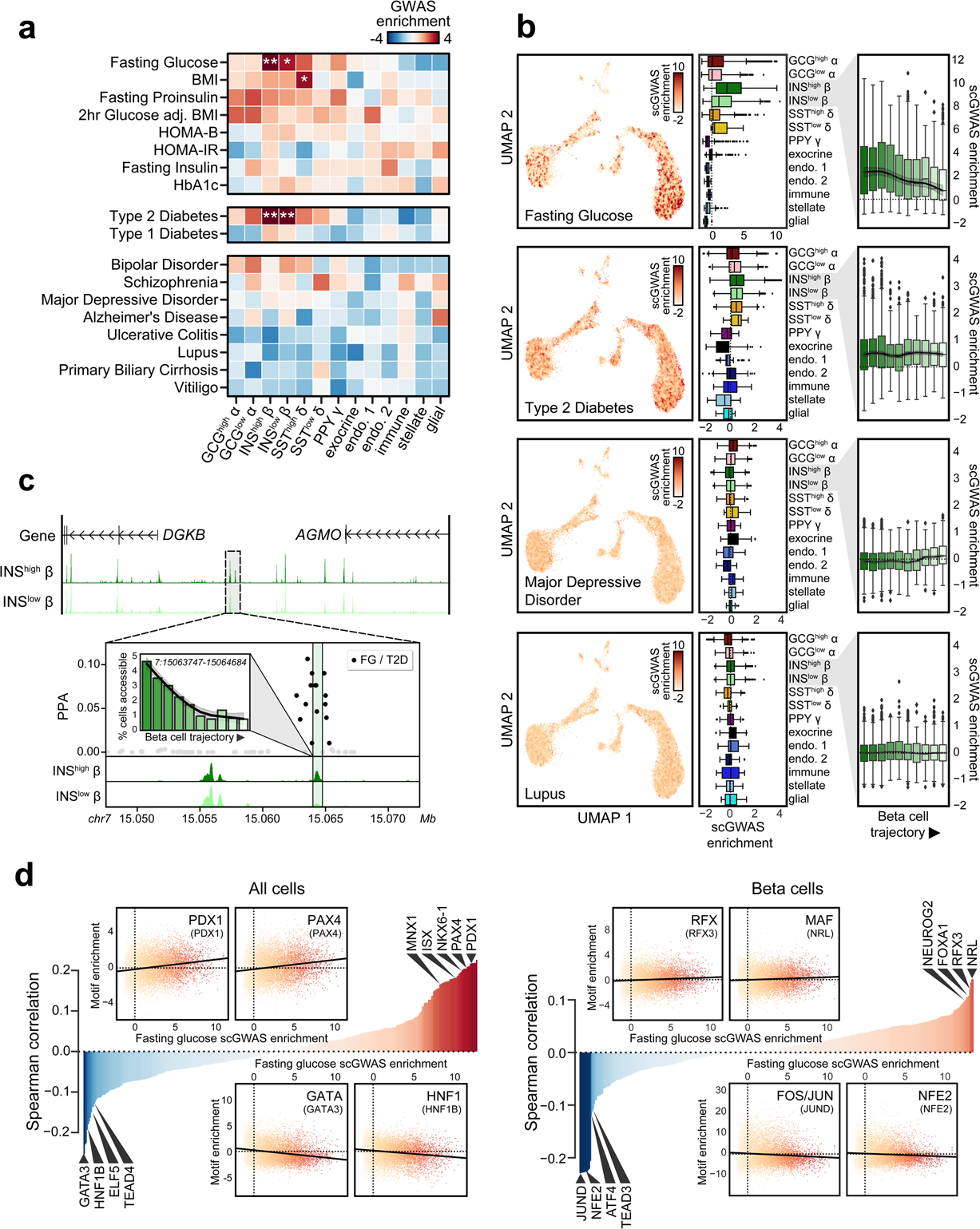
Enrichment of islet single cell accessible chromatin for diabetes and related trait genetic association data. (a) Cell type specific LD score regression enrichment z-scores for diabetes-related quantitative endophenotypes (top), type 1 and 2 diabetes (middle), and control traits (bottom) for islet snATAC-seq clusters. **FDR<.01 *FDR<.1. (b) Single cell enrichment z-scores for fasting glucose level, type 2 diabetes, major depressive disorder, and lupus projected onto UMAP coordinates (left panels), boxplot showing z-score enrichment distribution per cell type and state (middle panels), and z-score enrichment distribution split into 10 bins based on beta cell trajectory value (right panels). All boxplots show median (center line) and upper and lower quartiles (box limits). (c) Genome browser shot of the *DGKB* locus which is associated with both type 2 diabetes and fasting glucose level. Candidate causal variants fall in an enhancer with higher accessibility in INS^high^ beta cells and with dynamic chromatin accessibility decreasing across the beta cell trajectory, consistent with the beta cell enrichment patterns for fasting glucose level. (d) Correlation between single cell fasting glucose (FG) level enrichments and TF motif enrichments from chromVAR across all 14.2k cells (left) and 7.2k beta cells (right). Across all cells, FG level is positively correlated with beta cell TF motifs such as PDX1 and NKX6-1 and negatively correlated with alpha cell TF motifs such as GATA. Within beta cells, FG level is positively correlated with TF motifs enriched in the INS^high^ state such as RFX, NRL/MAF, and FOXA, and negatively correlated with TF motifs enriched in the INS^low^ state such as JUND and NFE2.

In these analyses, we again noted evidence for differences in enrichments between the hormone-high and -low states of endocrine cells (Figure 3a). To further resolve the heterogeneity of genetic association enrichment patterns, we used a novel framework to test the enrichment of genetic association signal genome-wide within accessible chromatin profiles of single cells (see Methods). We applied this approach to genetic association data for T2D and fasting glucose level, as well as negative control traits major depressive disorder and systemic lupus erythematosus (Figure 3b). We observed marked heterogeneity among beta cells in enrichment estimates for fasting glucose-associated variants, whereby cells in the INS^high^ state had significantly stronger enrichment than cells in the INS^low^ state (INS^high^ median Z=2.42, INS^low^ median Z=1.13, P<2.2×10^−16^) (Figure 3b). We further examined heterogeneity by calculating the average enrichment estimates for cells binned across the ‘pseudo-state’ trajectory (see Figure 2), which revealed a clear pattern of decreasing enrichment for fasting glucose-associated variation across pseudo-state moving from INS^high^ to INS^low^ beta cells (Figure 3b). Conversely, for T2D we observed enrichment for beta cells that was more consistent across INS^high^ and INS^low^ beta cells, as well as across the pseudo-state trajectory (INS^high^ median Z=0.48, INS^low^ median Z=0.51, P=0.84) (Figure 3b). In comparison, major depressive disorder and lupus showed no evidence for enrichment for beta cells (all median Z<.001) (Figure 3b). Knowledge of state-specific effects of cell types on specific phenotypes can then inform interpretation of association signals for those phenotypes; for example, at the *DGKB* locus, variants associated with both fasting glucose level and T2D overlapped a chromatin site with higher activity in INS^high^ beta cells, implicating this state-dependent regulatory sequence in mediating the association signal (Figure 3c).

Given our ability to map both complex trait and TF motif enrichments to single cells, we reasoned that joint analysis of these data could provide insights into TFs and regulatory networks through which genetic effects on these traits are mediated. We correlated single cell fasting glucose level and T2D enrichment z-scores with single cell TF motif enrichments from chromVAR^37^, both across all 14.2k islet cells as well as just the 7.2k beta cells (see Methods). Across all 14.2k cells, we observed strong positive correlation between fasting glucose level and T2D enrichment and beta cell lineage-specifying TF motifs (e.g. PDX1), and negative correlation with TF motifs regulating other islet cell types (Figure 3d, Supplementary Figure 9, Supplementary Table 6). When next considering only the 7.2k beta cells, we observed the strongest positive correlation between fasting glucose level and motifs in TF families enriched for INS^high^ beta cells (from Figure 2) such as RFX (ρ=.12, P=2.58×10^−24^), FOXA (ρ=.11, P=5.41×10^−19^), and MAF (ρ=.14, P=5.36×10^−32^), and negative correlation with INS^low^ beta cell TF motifs such as FOS/JUN and ATF (JUND ρ=-.23, P=1.23×10^−85^, ATF4 ρ=-.12, P=1.18×10^−23^) (Figure 3d, Supplementary Table 6). Interestingly, for T2D, both the strongest positive and negative correlations included motifs for TF families enriched in INS^low^ beta cell such as CCAAT and CREB (NFYA ρ=.073, P=1.72×10^−9^, CREB1 ρ=.053, P=7.44×10^−6^) and FOS/JUN (FOS::JUN ρ=-.06, P=2.45×10^−6^) (Supplementary Figure 9, Supplementary Table 6). Together these results provide state-resolved insight into the role of beta cells and beta cell TFs in T2D risk and fasting glucose level.

### Genome-wide predictions of variant effects on islet cell type- and state-specific regulatory sequence

Predicting the effects of non-coding genetic variants on regulatory activity remains a major challenge, in large part because the sequence vocabularies that encode regulatory function differ between cell types and states. Our cell type- and state-resolved accessible chromatin profiles provided an ideal opportunity to apply machine learning to model these regulatory vocabularies and use these models to predict the effects of genetic variants on putative regulatory sequences. We therefore used deltaSVM^70^ to predict the effects of genetic variants from the Haplotype Reference Consortium panel^71^ on chromatin accessibility in each endocrine cell type and cell state (see Methods). We identified 543,537 variants genome-wide with predicted allelic effects (FDR<.1), encompassing between 128k-210k variants (9.1%-14.8% of tested variants) per cell type or state (Figure 4a, Supplementary Data 3).

**Figure 4.**
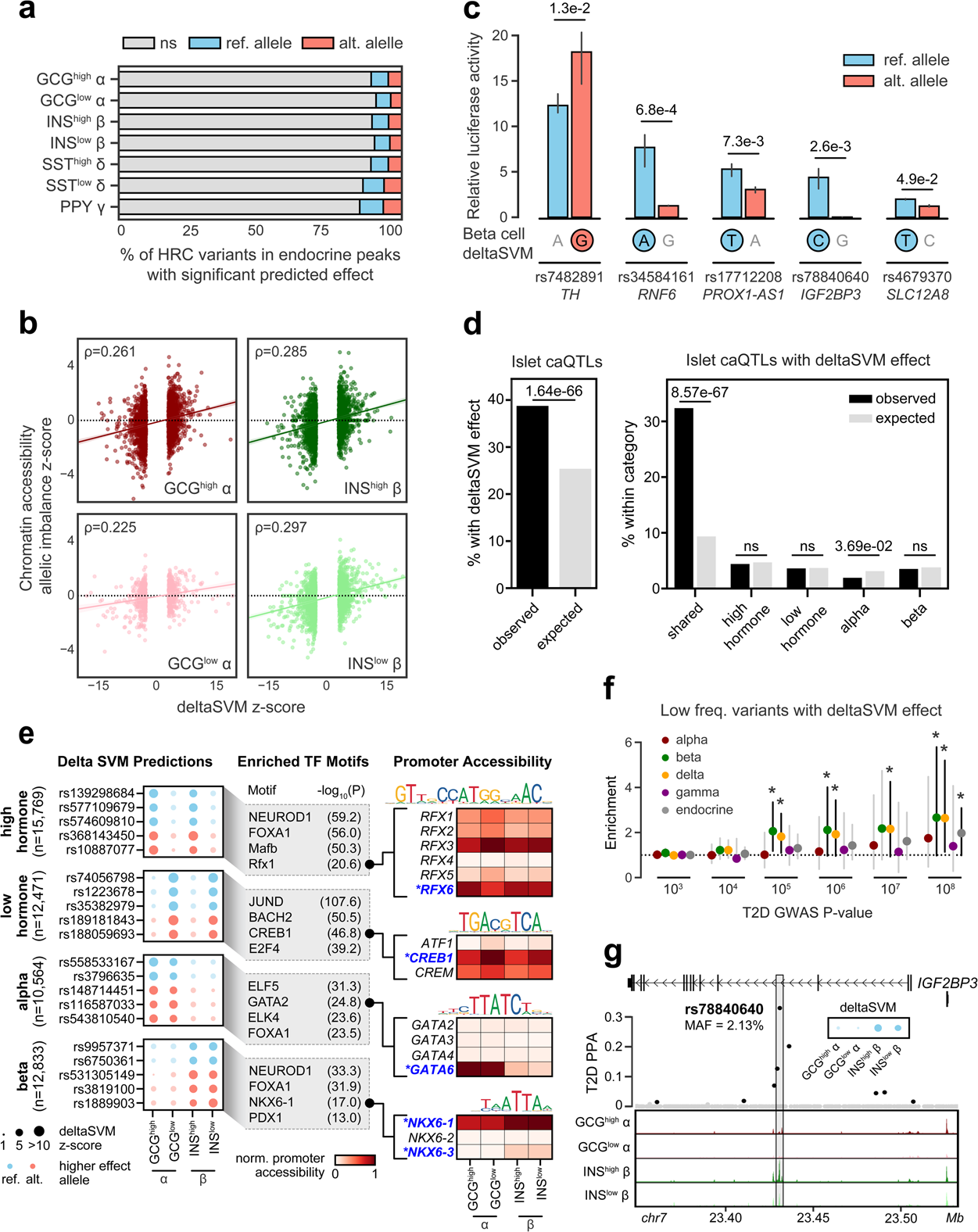
Genetic variants with islet cell type- and state-specific effects on chromatin accessibility. (a) Percentage of HRC reference panel r1.1 variants in any endocrine cell type peak (n=1,411,387 total) that had significant deltaSVM predictions at FDR<.1 for the reference (ref) or alternate (alt) allele in different endocrine cell types and states. (b) Spearman correlation comparing deltaSVM score to chromatin accessibility allelic imbalance z-scores using variants with significant deltaSVM predictions for alpha and beta states. (c) Luciferase gene reporter assays of five fine-mapped T2D variants with predicted beta cell effects in MIN6 cells. All tested variants (n=3) had significant effects in gene reporter assays and were directionally consistent with deltaSVM effects (highlighted with a circle around the allele with higher predicted effect). Data shown are mean ± 95% confidence interval. Two-sided Student’s T-test *P<.05 **P<.01 ***P<.001. (d) Enrichment of ensemble islet caQTLs for SNPs with significant deltaSVM effects in alpha and beta cells (left) and categorized based on shared, cell type- and state-specific deltaSVM effects on alpha and beta cells (right). Two-sided Fisher’s exact test. ns, not significant. (e) Variants with predicted cell type- and state-specific effects on alpha and beta cells, where size indicates magnitude of the deltaSVM z-score and color indicates the effect allele. Ref=blue, alt=red (left). TF motif families enriched in sequences surrounding the effect allele compared to the non-effect allele for each variant category (middle). Promoter accessibility patterns of genes in in enriched TF motif families. TFs with promoter accessibility patterns that match TF motif enrichment patterns are highlighted in blue and starred (right). (f) Enrichment of low frequency and rare variants with significant effects on islet chromatin for T2D association at different p-value thresholds. Data shown are enrichment ± 95% confidence interval. Two-sided binomial test *P<.05. (g) Low-frequency T2D-associated variant rs78840640 at the *IGF2BP3* signal has a high causal probability (PPA=0.33), overlaps peaks in both beta cell states, and is predicted to have allelic effects in beta cells.

To validate that our predictions captured true allelic effects on islet chromatin accessibility, we first compared alpha and beta cell predictions to allelic imbalance in chromatin accessibility measured directly from read count data at heterozygous variants in each sample (see Methods). We found significant correlations between predicted allelic effects and allelic imbalance estimates for all alpha and beta cell states (GCG^high^ Spearman ρ=.261, P=3.27×10^−46^, GCG^low^ ρ=.225, P=4.38×10^−10^, INS^high^ ρ=.285, P=1.13×10^−53^, INS^low^ ρ=.297, P=2.28×10^−40^) (Figure 4b). We further validated five likely causal T2D variants identified in fine-mapping studies and predicted to have allelic effects on beta cell chromatin using gene reporter assays in the MIN6 beta cell line. In each case, reporter assays showed significant allelic effects on enhancer activity that were directionally consistent with predictions (Figure 4c). We also compared predictions to chromatin accessibility quantitative trait loci (caQTLs) previously identified in ensemble islet samples^72^. We observed highly significant enrichment of caQTLs among variants with predicted effects on alpha or beta cells (obs.=38.8%, exp.=23.6%, two-sided Fisher’s exact P=1.64×10^−66^) (Figure 4d). When sub-dividing predictions based on those with shared, cell type-specific (alpha, beta) or state-specific (hormone-high, hormone-low) effects we observed significant enrichment of caQTLs only among shared effect variants (Figure 4d), suggesting that islet caQTLs may have lower sensitivity for variants with cell type- or state-specific effects.

We thus sought to further characterize genetic variants predicted to have cell type- and state-dependent effects on islet chromatin. For each category of variants, we performed motif enrichment comparing sequences around the effect allele to the non-effect allele (see Methods). Variants with state-specific effects tended to disrupt motifs for TF families such as NEUROD, FOXA, MAF and RFX for hormone-high states (-log_10_(P)=59.2, 56.0, 50.3, 20.6), and signaling-responsive TF families such as JUN/FOS and CREB for hormone-low states (-log_10_(P)=107.6, 46.8) (Figure 4e). Similarly, variants with alpha or beta cell-specific effects tended to disrupt motifs for lineage-defining TFs and TF families including GATA for alpha cells (-log_10_(P)=24.8), and NKX6 and PDX1 for beta cells (-log_10_(P)=17.0, 13.0) (Figure 4e). In order to assign motifs to specific TFs, we again examined promoter-accessibility of TFs within the structural TF subfamily^48^ (see Methods). For example, among GATA subfamily members only GATA6 had high promoter accessibility in alpha cells (GCG^high^: 1.00, GCG^low^: .97, INS^high^: .21, INS^low^: .13), suggesting that GATA6 binding is likely disrupted in alpha cells by variants affecting the GATA motif. Similarly, among NKX6 subfamily members, NKX6-1 and NKX6-3 had promoter accessibility in beta cells (NKX6-1 GCG^high^: .78, GCG^low^: .80, INS^high^: .98, INS^low^: 1.00; NKX6-3 GCG^high^: 0, GCG^low^: 0, INS^high^: .18, INS^low^: .19), and among RFX family members RFX6 had promoter accessibility in hormone-high state cells (GCG^high^: .93, GCG^low^: .68, INS^high^: 0.88, INS^low^: .85) (Figure 4e).

Predictions of allelic effects are particularly important in interpreting the function of low frequency non-coding variants, which are impractical to assay by standard approaches such as QTL mapping without very large sample sizes. We thus evaluated whether our predictions could prioritize lower frequency (defined as minor allele frequency [MAF]<.05) functional variants involved in T2D risk. We compared T2D association at different p-value thresholds for lower frequency variants with significant effects for any endocrine cell type, as well as for each cell type individually, to background variants without predicted effects (see Methods). We observed enrichment of genome-wide significant T2D associations among lower frequency variants with predicted effects in any endocrine cell type compared to background (Figure 4f). When considering effects in each cell type, we observed enrichment of T2D association among variants with predicted effects in beta cells as well as delta cells, even down to sub-genome-wide significant p-values (Figure 4f). We next highlighted specific low frequency, T2D risk variants with predicted effects. At the *IGF2BP3* locus, rs78840640 (MAF=.02) had allelic effects on beta cell chromatin (INS^high^ beta q=.0015; INS^low^ beta q=.041), and fine-mapping data supported a causal role in T2D (posterior probability [PPA]=.33) (Figure 4g). We confirmed in gene reporter assays that this variant affected enhancer activity where the alternate (and T2D risk) allele G had reduced activity (Figure 4c). We also observed predicted effects for rare T2D variants for example rs186384225 (MAF=.0037) at *TCF7L2* and rs571342427 (MAF=.0015) at *INS-IGF2* (Supplementary Figure 10a-b). These results reveal that cell type-specific chromatin can provide accurate functional predictions of lower frequency variants, enabling more effective interpretation of genome sequence from patients and disease cohorts.

### Co-accessibility links distal regulatory variants to putative target genes in distinct islet cell types and states

Defining the genes affected by regulatory element activity remains a major challenge, as enhancers can regulate gene activity over large, non-adjacent distances^73^. A number of approaches have been developed to link regulatory elements to target genes including 3D chromatin architecture assays and correlation of accessible chromatin activity across multiple samples^74,75^. While these approaches have different strengths, a common weakness is reliance on ensemble data and non-cell type-resolved information^27,76^. Recently, a new approach was developed to link regulatory elements based on co-accessibility across single cells^6^, which has the potential to provide cell type-resolved enhancer-promoter relationships. We thus sought to leverage accessible chromatin profiles across thousands of islet cells to define co-accessibility between sites in specific cell types. For these analyses we again focused on alpha and beta cells where cell numbers (5,594 and 7,170 cells, respectively) gave us the most power to effectively derive co-accessibility maps.

To calibrate the extent to which co-accessibility reflected physical interactions between regulatory elements, we first performed a distance-matched comparison between co-accessible sites stratified by co-accessibility threshold to chromatin loops identified from Hi-C and promoter capture Hi-C (pcHi-C) assays in primary islets^27,76^. We observed strong enrichment for pairs of sites with co-accessibility scores >.05 in both alpha and beta cells for islet chromatin loops identified from pcHi-C and Hi-C compared to sites that had no evidence for co-accessibility (Figure 5a, Supplementary Figure 11a-c). We therefore used this threshold (.05) to define co-accessibility, through which we identified 593,769 co-accessible sites in alpha cells (Supplementary Data 4) and 487,549 co-accessible sites in beta cells (Supplementary Data 5). There were 64,045 (alpha) and 57,374 (beta) unique distal sites co-accessible with a gene promoter (median 2 promoters per site), and 19,872 (alpha) and 19,269 (beta) unique gene promoters co-accessible with a distal site (median 9 per gene in alpha, 6 in beta cells) (Supplementary Figure 11d-e).

**Figure 5.**
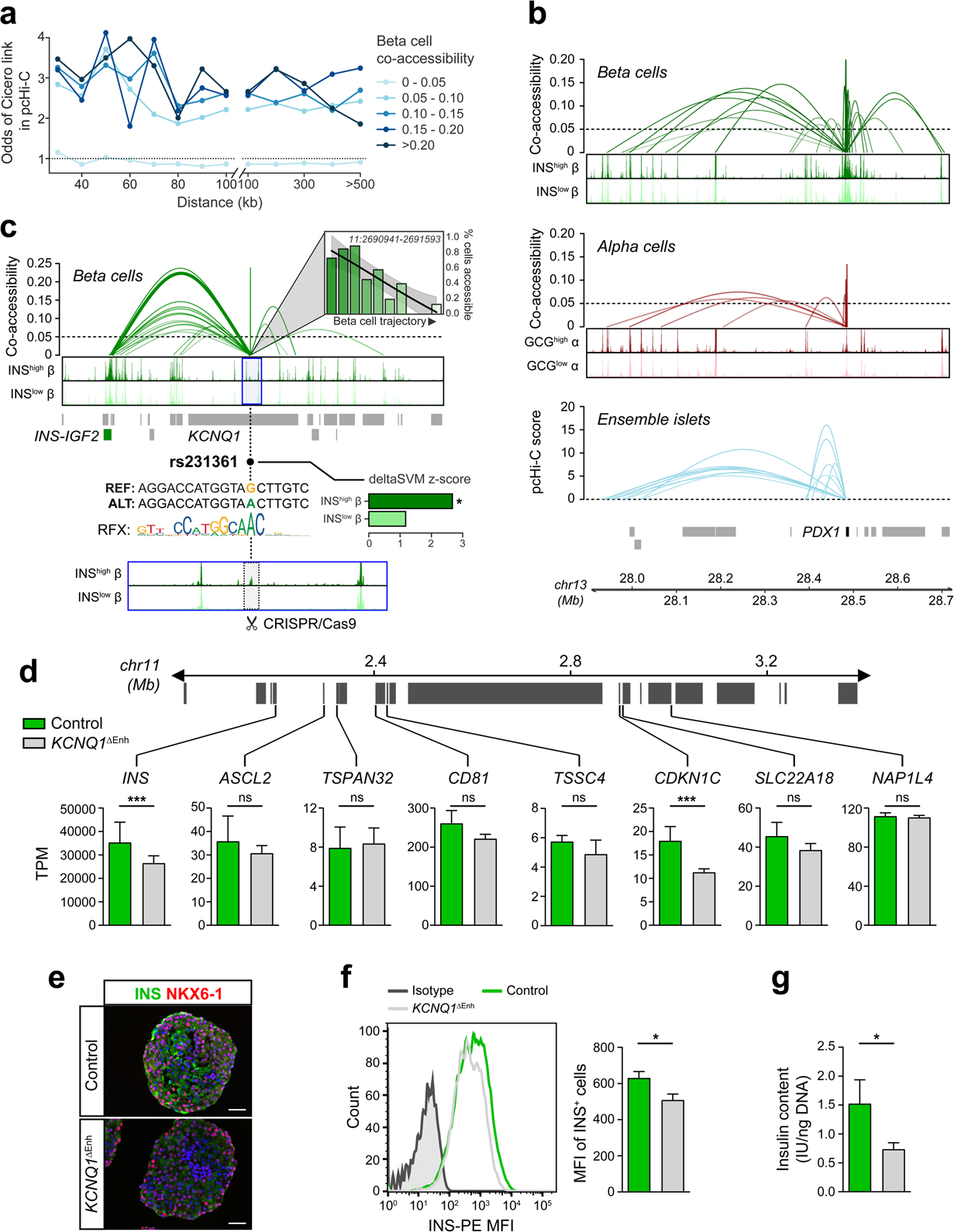
Chromatin co-accessibility links cell type enhancers and diabetes risk variants to target genes. (a) Distance-matched odds that beta cell co-accessibility links overlap islet pcHi-C chromatin loops at different co-accessibility threshold bins. (b) Beta cell (top) and alpha cell (middle) co-accessibility between pairs of accessible chromatin sites and high-confidence promoter capture Hi-C interactions from bulk islets (bottom) anchored at the *PDX1* promoter. (c) Beta cell co-accessibility anchored on an enhancer within *KCNQ1* harboring causal T2D variant rs231361 (PPA=1) shows distal links to the *INS* promoter as well as other non-promoter sites. This enhancer has an accessible peak call in the INS^high^ beta cell state but not the INS^low^ state and has dynamic accessibility across the beta cell state trajectory. rs231361 disrupts a sequence motif for *RFX,* which itself is enriched in INS^high^ beta cells, has dynamic enrichment across the beta cell trajectory, and is predicted to have allelic effects on INS^high^ beta cells (deltaSVM z-score *FDR<.1). We performed CRISPR/Cas9-mediated deletion of the 2.6 kb genomic region flanking this enhancer (highlighted in grey) in hESCs (*KCNQ1*^ΔEnh^). (d) Differential expression analysis of genes within 2 Mb of the *KCNQ1* enhancer in beta cell stage cultures (day 28) from *KCNQ1*^ΔEnh^ (n=6; 3 clones each differentiated two times) and control (n=2; 1 clone differentiated two times) hESC clones. *INS* and *CDKN1C* mRNA levels are significantly reduced in *KCNQ1*^ΔEnh^ compared to control cells, while other genes in the region show no significant difference in expression. Data are shown as transcripts per million (TPM). (e) Representative immunofluorescence staining for INS (green) and NKX6-1 (red) with DAPI staining (blue) on beta cell stage *KCNQ1*^ΔEnh^ and control aggregates. Scale bar, 50μm. (f) Histogram showing INS fluorescence intensity by flow cytometry (left panel) and quantification of INS median fluorescence intensity (MFI, right panel) in beta cell stage cultures from *KCNQ1*^ΔEnh^ (n=9; 3 clones each differentiated three times) and control (n=6; 2 clones each differentiated three times) cells. (g) Insulin content in beta cell stage cultures from *KCNQ1*^ΔEnh^ (n=9; 3 clones each differentiated three times) and control (n=6; 2 clones differentiated three times) clones. Data are shown as mean ± SEM. * p < 0.05, *** p<0.001, ns, not significant by two-sided Student’s T-test.

Among co-accessible links to gene promoters, the majority (71.9%) were alpha or beta cell-specific, highlighting the value of single cell-resolved data for identifying putative cell type-specific regulatory interactions. As an example of cell type-specific co-accessibility, the *PDX1* promoter had co-accessibility with 35 sites in beta cells, including a site over 500 kb distal that directly coincided with an islet pcHi-C loop, only 7 of which were also found in alpha cells (Figure 5b). In another example, at the *ARX* locus, 17 sites were co-accessible with the *ARX* promoter in alpha cells, none of which were co-accessible in beta cells (Supplementary Figure 11f). Conversely, as an example of shared co-accessibility across cell types, the *NEUROD1* promoter was co-accessible with 52 and 47 chromatin sites in alpha and beta cells, respectively, of which 26 were shared and several were over 500 kb distal (Supplementary Figure 11g).

Given heterogeneity in alpha and beta cell regulatory programs, we next cataloged co-accessible links between distal alpha and beta cell sites and gene promoters that had differential activity across and hormone-high and -low states (see Methods). We observed 25,012 (alpha) and 9,641 (beta) co-accessible links where both the distal site (unique distal sites: alpha=10,926, beta=7,958) and the gene promoter (unique promoters: alpha=1,951, beta=1,516) were differentially active between states in the same direction. State-dependent co-accessible links included both gene promoters active in the hormone-high state such as *INS*, *GCG, G6PC2,* and *NEUROD1*, and gene promoters active in the hormone-low state such as *FOSL1, FOSL2, CREB1,* and *CREB5*. We also identified genes with different isoform promoters co-accessible with hormone-high and hormone-low dependent distal sites such as *GLIS3* (Supplementary Figure 11h), suggesting these genes have distinct regulatory programs driving isoform-specific activity across different cell states.

Distal sites with co-accessibility links to gene promoters harbored risk variants for T2D at many loci, suggesting this approach can prioritize target genes of T2D risk variants in islets. We observed one such example at the *KCNQ1* locus, where an islet chromatin site located in intron 3 of *KCNQ1* had beta cell-specific co-accessibility with the *INS* promoter over 500 kb distal and harbored a causal T2D risk variant rs231361 (PPA=1)^10^. (Figure 5c). Published 4C data from the EndoC-βH1 human beta cell line^77^ anchored on the *INS* promoter supported the existence of physical interactions between this site and the *INS* promoter in beta cells (Supplementary Figure 12a). Interestingly, the site was more accessible in INS^high^ beta cells compared to INS^low^ beta cells, and rs231361 was predicted to have state-specific effects on beta cell chromatin accessibility (INS^high^ beta FDR q=.060; INS^low^ beta FDR q=.40). Furthermore, rs231361 disrupted an *RFX* family sequence motif, which itself was enriched in the INS^high^ beta cell state (Figure 5c, also see Figure 2c). The *KCNQ1* locus is also associated with quantitative measures of insulin secretion^78–81^ and fasting glucose level^82^, suggesting that the mechanism of action of this locus on T2D risk is likely mediated through beta cell function in a state-dependent manner.

To validate the effects of the chromatin site containing rs231361 on distal regulation of *INS* in beta cells, we deleted a 2.6 kb region flanking the site in human embryonic stem cells (hESCs) by CRISPR/Cas9-mediated genome editing, generating three bi-allelic deletion clones (*KCNQ1*^ΔEnh^) (Figure 5c, Supplementary Figure 12b-c). We then differentiated the three *KCNQ1*^ΔEnh^ clones as well as two unedited control clones into beta cells using an established protocol^83^ with minor modifications (see Methods). Analysis of cultures at the beta cell stage revealed similar numbers of INS^+^ cells in *KCNQ1*^ΔEnh^ and control clones (91.1±4.02% vs 94.6±2.11%) (Supplementary Figure 12d), suggesting that the enhancer deletion had no effect on beta cell differentiation. Further supporting this conclusion, similar numbers of cells expressed the beta cell marker NKX6-1 in *KCNQ1*^ΔEnh^ and control cultures (Supplementary Figure 12e). Likewise, *NKX6-1* mRNA levels were similar (FDR=0.98) (Supplementary Figure 12f). Next, we determined effects of the enhancer deletion on gene expression in *cis,* interrogating all genes within 2 Mb of the enhancer. We observed a significant decrease in the expression of *INS* (P=3.02×10^−4^; FDR=0.066) and *CDKN1C* (P=1.96×10^−4^; FDR=0.059) in *KCNQ1*^ΔEnh^ compared to control cells, whereas other genes in the region showed no difference in expression (all P>.05; note *KCNQ1* itself was not expressed) (Figure 5d). Analysis of INS protein by immunofluorescence staining, flow cytometry, and ELISA further revealed reduced INS protein abundance in *KCNQ1*^ΔEnh^ beta cells (Figure 5e-g). In contrast, beta cell NKX6-1 protein levels were not affected (Supplementary Figure 12e), confirming specific effects of the enhancer deletion on *INS* mRNA and protein expression in beta cells.

### A resource of islet cell type and state regulatory programs to annotate T2D risk variants

Together our results provide a multi-tiered reference of islet cell type and cell state regulatory programs through which non-coding genetic variants can be comprehensively annotated. As most genetic risk variants for diabetes are non-coding, this resource can be used to interpret their molecular mechanisms. We therefore annotated the islet cell type-specific regulatory programs of T2D risk variants using fine-mapping ‘credible sets’ of 402 risk signals^10,84^. Fine-mapped credible set variants at 239 risk signals mapped within an islet cell type chromatin site and, at 97 of these 239 risk signals, credible set variants also had both predicted allelic effects and co-accessibility with a gene promoter (Supplementary Table 7).

Genes co-accessible with fine-mapped credible set T2D variants in islet cell type chromatin were enriched for biological processes related to protein localization and transport, stress response, cell cycle, and signal transduction (Supplementary Table 8). Co-accessible genes also included numerous genes involved in monogenic diabetes such as *INS, KCNJ11, ABCC8, HNF1A, HNF4A, GCK,* and *NKX2-2*, as well as TFs in structural families with lineage- and state-specific motif enrichments (from Figures 1-2) such as *NKX6-1, NFATC2* and *RFX6*. At 22 T2D loci, fine-mapped variants at multiple independent risk signals were co-accessible with the same gene, providing independent support for the role of these genes in T2D. For example, at the *KCNQ1* locus (11p15), fine-mapped variants at four T2D risk signals (including rs231361 above) were in sites co-accessible with the *INS* promoter (Supplementary Figure 13a), and at the *CDKN2A/B* locus (9p21), fine-mapped variants at five T2D signals were in sites co-accessible with the *CDKN2A, MTAP* and *DMRTA1* promoters (Supplementary Figure 13b). In other examples, at the *DGKB* locus (7p21), fine-mapped variants at two T2D signals were in sites co-accessible with the *DGKB* promoter (Supplementary Figure 13c), and at 7p13 fine-mapped variants at two T2D signals were in sites co-accessible with the *GCK* promoter (Supplementary Figure 13d).

In order to effectively provide these data to the community to facilitate hypothesis testing and mechanistic discovery, we developed a publicly-accessible web portal and database (https://www.t2depigenome.org) which contains processed data and islet cell type annotations generated in this study, as well as epigenomic data from islets and other diabetes-relevant tissues available in other published studies (Supplementary Figure 14a-f). In addition, the portal enables the user to query genetic variants for their respective islet cell type annotations.

## Discussion

Our findings provide a roadmap demonstrating how single cell accessible chromatin data derived from disease-relevant primary tissue can be utilized to define the cell types, cell states, *cis* regulatory elements and genes involved in the genetic basis of complex disease. Over 400 known risk signals for T2D have been identified, yet only a handful have been characterized molecularly^16,18,27, 85–91^. Identifying the genes affected by non-coding risk variants is paramount for understanding the molecular pathways dysregulated in disease and can inform therapeutic target discovery. Candidate target genes of T2D risk signals derived using single cell co-accessibility were highly enriched for disease-relevant biological processes, and many of these genes serve as compelling targets for mechanistic study. At the *KCNQ1* locus, co-accessibility data and hESC beta cell models revealed that a long-range enhancer harboring a causal T2D variant affects insulin expression and protein levels in beta cells. Mutations of *INS* cause monogenic diabetes and tandem repeats in *INS* affect T1D risk^92,93^, but to our knowledge *INS* has not been directly implicated in T2D risk. The *KCNQ1* locus has a complex contribution to T2D with 10 signals in the region that each confer independent risk^10^, four of which had beta cell co-accessibility with the *INS* promoter. We therefore speculate that the *KCNQ1* locus mediates T2D risk through multiple long-range regulatory effects on *INS*, in addition to *CDKN1C* and other genes.

Single cell accessible chromatin uncovered heterogeneity in the regulatory programs of endocrine cell types, revealing cell type- and state-resolved effects of genetic variants on fasting glucose and T2D risk. Previous studies have characterized heterogeneity in beta cell physiological function, cell surface markers, and gene expression^22,94,95^. The heterogeneity we observed in the beta cell epigenome mapped to cellular states related to insulin production and stress-related signaling response^23^, and we identified TFs likely driving cell state-specific functions. Integrating single cell heterogeneity with large-scale genetic association data revealed that genetic variants modulating fasting glucose levels likely act through the insulin-producing beta cell state, whereas genetic risk of T2D is mediated through both the insulin-producing state and other functional beta cell state(s). Moreover, given similar heterogeneity in the epigenomes of alpha and delta cells, our results reveal that endocrine cell regulation involves both lineage-specific programs as well as an additional layer of state-specific programs common across endocrine cell types.

In summary, we present the most detailed characterization of islet cell type and state regulatory programs to date and a web resource to query these programs. When combined with genetic fine-mapping and genome sequencing, this resource will greatly enhance efforts to define molecular mechanisms of T2D risk. More broadly, our study provides a framework for using single cell chromatin from disease-relevant tissues to interpret the genetics and biological mechanisms of complex disease.

## Methods

### Islet processing and nuclei isolation

We obtained islet preparations from three donors for the Integrated Islet Distribution Program (IIDP) (Supplementary Table 1). Islet preparations were further enriched using zinc-dithizone staining followed by hand picking. Studies were given exempt status by the Institutional Review Board (IRB) of the University of California San Diego.

### Generation of snATAC-seq libraries

Combinatorial barcoding single nuclear ATAC-seq was performed as described previously^2,4^ with several modifications as described below. For each donor (N=3), approximately 3,000 islet equivalents (IEQ, roughly 1,000 cells each) were resuspended in 1 ml nuclei permeabilization buffer (10mM Tris-HCL (pH 7.5), 10mM NaCl, 3mM MgCl_2_, 0.1% Tween-20 (Sigma), 0.1% IGEPAL-CA630 (Sigma) and 0.01% Digitonin (Promega) in water) and homogenized using 1ml glass dounce homogenizer with a tight-fitting pestle for 15 strokes. Homogenized islets were incubated for 10 min at 4°C and filtered with 30 µm filter (CellTrics). Nuclei were pelleted with a swinging bucket centrifuge (500 x g, 5 min, 4°C; 5920R, Eppendorf) and resuspended in 500 µL high salt tagmentation buffer (36.3 mM Tris-acetate (pH = 7.8), 72.6 mM potassium-acetate, 11 mM Mg-acetate, 17.6% DMF) and counted using a hemocytometer. Concentration was adjusted to 4500 nuclei/9 µl, and 4,500 nuclei were dispensed into each well of a 96-well plate. Glycerol was added to the leftover nuclei suspension for a final concentration of 25 % and nuclei were stored at −80°C. For tagmentation, 1 µL barcoded Tn5 transposomes^4,96^ were added using a BenchSmart™ 96 (Mettler Toledo), mixed five times and incubated for 60 min at 37 °C with shaking (500 rpm). To inhibit the Tn5 reaction, 10 µL of 40 mM EDTA were added to each well with a BenchSmart™ 96 (Mettler Toledo) and the plate was incubated at 37 °C for 15 min with shaking (500 rpm). Next, 20 µL 2 x sort buffer (2 % BSA, 2 mM EDTA in PBS) were added using a BenchSmart™ 96 (Mettler Toledo). All wells were combined into a FACS tube and stained with 3 µM Draq7 (Cell Signaling). Using a SH800 (Sony), 20 nuclei were sorted per well into eight 96-well plates (total of 768 wells) containing 10.5 µL EB (25 pmol primer i7, 25 pmol primer i5, 200 ng BSA (Sigma), PMID: 29434377). Preparation of sort plates and all downstream pipetting steps were performed on a Biomek i7 Automated Workstation (Beckman Coulter). After addition of 1 µL 0.2% SDS, samples were incubated at 55 °C for 7 min with shaking (500 rpm). We added 1 µL 12.5% Triton-X to each well to quench the SDS and 12.5 µL NEBNext High-Fidelity 2× PCR Master Mix (NEB). Samples were PCR-amplified (72 °C 5 min, 98 °C 30 s, (98 °C 10 s, 63 °C 30 s, 72 °C 60 s) × 12 cycles, held at 12 °C). After PCR, all wells were combined. Libraries were purified according to the MinElute PCR Purification Kit manual (Qiagen) using a vacuum manifold (QIAvac 24 plus, Qiagen) and size selection was performed with SPRI Beads (Beckmann Coulter, 0.55x and 1.5x). Libraries were purified one more time with SPRI Beads (Beckmann Coulter, 1.5x). Libraries were quantified using a Qubit fluorimeter (Life technologies) and the nucleosomal pattern was verified using a Tapestation (High Sensitivity D1000, Agilent). The library was sequenced on a HiSeq2500 sequencer (Illumina) using custom sequencing primers, 25% spike-in library and following read lengths: 50 + 43 + 40 + 50 (Read1 + Index1 + Index2 + Read2).

### Raw data processing and quality control

For each read, we first appended the cell barcode metadata to the read name. The cell barcode consisted of four pieces (P7, I7, I5, P5) which were derived from the index read files. We first corrected for sequencing errors by calculating the Levenshtein distance between each of the four pieces and a whitelist of possible sequences. If the piece did not perfectly match a whitelisted sequence, we took the best matching sequence if it was within 2 edits and the next matching sequence was at least 2 additional edits away. If none of these conditions were met, we discarded the read from further analyses.

We trimmed Nextera adapter sequences from sequence reads using trim_galore (v.0.4.4, https://github.com/FelixKrueger/TrimGalore) with default parameters. We used bwa mem^97^ (v.0.7.17-r1188) to align reads to the hg19 reference genome with the options ‘-M -C’. We then used samtools^98^ to filter out reads that did not align to the autosomes or sex chromosomes and low mapping quality reads (MAPQ<30). We used samtools fixmate (v.1.6) to perform additional checks for FR proper pairs and removed secondary or unmapped reads. We used the MarkDuplicates tool from picard (https://broadinstitute.github.io/picard/) to remove duplicates on a per-barcode basis with ‘BARCODE_TAG’ option. For each experiment, we used a Gaussian mixture model on log-transformed read depths to separate barcodes with a 99% probability of belonging to the high read distribution, likely representing real cells, from those in the low read distribution, likely representing background reads. We then set an additional threshold of 1000 read depth, reasoning that low read cells would contribute additional noise to clustering.

### Cluster analysis for snATAC-seq

We split the genome into 5 kb windows and removed windows overlapping blacklisted regions from ENCODE (https://sites.google.com/site/anshulkundaje/projects/blacklists). For each experiment, we then created a sparse *m* x *n* matrix containing read depth for *m* cells passing read depth thresholds at *n* windows. For further quality checks, we performed initial clustering for each experiment individually using scanpy^99^ (v.1.4). We extracted highly variable windows using mean read depths and normalized dispersion. After normalization to a uniform read depth and log-transformation of read depth, we regressed out the log-transformed total read depth for each cell. We then performed PCA and extracted the top 50 principal components. We used these components to calculate the nearest 30 neighbors using the cosine metric, which were subsequently used for UMAP dimensionality reduction with the parameters ‘min_dist=0.3’ and Louvain clustering with the parameters ‘resolution=1.5’. For each experiment, we removed 2,709 cells that were in clusters corresponding to low read depth.

After removing these cells, we used similar methods to cluster cells from all experiments together with the following modifications. We extracted highly variable windows across cells from all experiments. Since read depth was a technical covariate specific to each experiment, we regressed this out on a per-experiment basis. We used mutual nearest neighbors correction^30^ (mnnpy, v.0.1.9.4) to adjust for batch effects across experiments with the parameters ‘k=10’. We then performed clustering as described above. We used chromatin accessibility at windows overlapping promoters for marker hormones (*GCG*, *INS-IGF2*, *SST*, and *PPY*) to assign cell types for the endocrine islet cell types (alpha, beta, delta, and gamma). We performed re-clustering on non-endocrine islet clusters and used chromatin accessibility at windows around marker genes from single cell RNA-seq to assign cluster labels. In our clustering results, we identified a cluster of 694 alpha cells that were mostly derived from a single donor (96% of cells from Islet 1). Because we were uncertain whether this represented technical or biological differences, we excluded this cluster from further analyses. We also excluded a cluster of 192 cells likely representing lower quality cells as it had low intra-cluster similarity and lower fraction of reads in peaks.

### Comparison to bulk and sorted islet ATAC-seq

We obtained raw sequence data of ATAC-seq for 42 bulk islet samples from four prior studies^14,27,28,72^ and 4 bulk pancreas samples from ENCODE. We re-processed all samples with a uniform pipeline: we aligned all reads to hg19 with bwa mem, identified and removed duplicate reads with picard MarkDuplicates, and called peaks with MACS2 (v.2.1.2) with the parameters ‘—shift −100 –extsize 200 –keep-dup all’. For the three islet snATAC-seq samples, we used aggregated per-barcode deduplicated reads to call peaks. We defined all possible accessibility peaks by filtering out ENCODE blacklisted regions and retaining merged peaks on autosomal chromosomes found in more than one sample. We then calculated the read coverage at all possible accessibility peaks and TPM-normalized the counts. We calculated the Spearman correlation between normalized read coverages and used hierarchical clustering to assess similarity between bulk islet samples. To check peak call overlap between aggregated single cell ATAC and bulk ATAC data, we split peaks based into promoter proximal (+/-500 bp from GENCODE transcript TSS) and distal peaks based on promoter overlap. For each cluster, we calculated the percentage of aggregate peaks that overlapped merged autosomal bulk peaks and individual sample-level autosomal bulk peaks.

We also obtained raw sequence data of ATAC-seq from flow-sorted pancreatic cells (alpha, beta, acinar, ductal) from two prior studies^35,36^ and re-processed all samples with the uniform pipeline described above. For alpha, beta, and exocrine cells from islet snATAC-seq, we split reads on a per-donor and per-cluster basis to obtain read files. Because total read depth was highly variable across sorted samples, we merged autosomal peaks after filtering out ENCODE blacklist regions. We calculated read coverage in each sample for each merged peak and TPM normalized count values. We then calculated the Spearman correlation between normalized read coverages and used hierarchical clustering to assess similarity between sorted and snATAC-seq islet samples.

### Identifying marker peaks of chromatin accessibility

To identify peaks for each cell type, we aggregated reads for all cells within a cluster or sub-cluster. We shifted reads aligning to the positive strand by +4 bp and reads aligning to the negative strand by −5 bp, extended reads to 200 bp, and centered reads. We used MACS2^100^ to call peaks of chromatin accessibility for each aggregated read file with the following settings ‘--nomodel -- keep-dup all’. We removed peaks that overlapped ENCODE blacklisted regions^101^. We then used bedtools^102^ to merge peaks from all clusters and sub-clusters to create a superset of islet regulatory peaks.

We generated a sparse *m* x *n* binary matrix containing binary overlap between *m* peaks in the superset of islet regulatory peaks and *n* cells. We then calculated t-statistics of peak specificity for each cluster or sub-cluster through linear regression models. We used binary encodings to specify which donor a given cell came from as covariates in the model. For each peak and cluster, we used binary encoding of read overlap with the peak as the predictor and whether a cell was in the cluster (1 if yes, −1 if no) as the outcome.

### Matching islet snATAC-seq with scRNA-seq clusters

To verify that clusters definitions and labels from single cell chromatin accessibility data matched those from single cell expression data, we obtained published single cell RNA-seq data from 12 non-diabetic islet donors^23^. Because cluster definitions for all cell types were not available, we re-analyzed the data and performed clustering analyses. Starting with the gene expression matrix, we first performed quality control steps to remove potential doublets. For each marker gene of different cell types *GCG* (alpha), *INS* (beta), *SST* (delta), *PPY* (gamma), *CTRB2* (acinar), *CFTR* (ductal)*, PLVAP* (endothelial)*, PDGFRB* (stellate), and *C1QC* (immune) we used a Gaussian mixture model on log-transformed read depth to determine whether a cell expressed the gene (high distribution) or not (low distribution). We verified that cells expressing more than one marker gene had on average higher read depth and expressed more genes (Supplementary Figure 3a,b). We regressed out covariates including sex, BMI, and read depth, and separated cells by donor of origin. We then used MNN correction^30^ to adjust for batch effects. After scaling the data, we performed PCA and used the top 50 principal components to calculate the 10 nearest neighbors using the cosine metric. We used the nearest neighbor map for UMAP dimensionality reduction with the parameters ‘min_dist=0.3’ and to perform Louvain clustering with the parameters ‘resolution=1’ (Supplementary Figure 3c). We used a similar regression framework as the chromatin accessibility marker peaks to calculate t-statistics for gene specificity for each cluster (Supplementary Figure 3d,e) with the following modifications: we included sex, BMI, and log-transformed read coverage as covariates and used log_2_ read counts for each gene instead of binary peak coverage as the predictor.

We used the Spearman correlation between t-statistics from islet snATAC-seq and scRNA-seq data to match up clusters. Specifically, we took the top 100 (sorted by descending t-statistic) most specific promoter peaks for each cluster or sub-cluster to define a list of genes for comparison. To facilitate one-to-one comparisons between the two datasets, for this analysis only we defined promoter peaks as peaks within +/-500 bp of a GENCODE v19 gene TSS. This list contained 966 genes, which is less than 100×13 (number of clusters) because 1) marker genes were sometimes shared between sub-clusters and 2) not all genes were present in the expression dataset. For each cluster from accessible chromatin data, we then compared t-statistics of genes in the list with t-statistics for all clusters from single cell expression using the Spearman correlation, which is robust to very specific marker genes such as insulin which could otherwise bias these comparisons.

### Motif enrichment with chromVAR

We used chromVAR^37^ (v.1.5.0) to calculate TF motif-associated difference between cell populations. We first calculated counts per peak per cell matrix and then input it to chromVAR. We filtered cells with minimal reads less than 1500 (min_depth=1500) and peaks with fraction of reads less than 0.15 (min_in_peaks=0.15) by using ‘filterSamplesPlot’ function from chromVAR. We also corrected GC bias based on ‘BSgenome.Hsapiens.UCSC.hg19’ using ‘addGCBias’ function. Then we used the Jaspar motifs from ‘getJasparMotifs’ function with default parameter and calculated the deviation z-scores for each TF motif in each cell by using ‘computeDeviations’ function. High-variance TF motifs across all cell types were selected by ‘computeVariability’ function using cut-off 1.2 (N=111). For each of these variable motifs, we calculated the mean z-score for each cell types and normalized the values to 0 (minimal) and 1 (maximal).

### Comparison of alpha and beta cell states

To identify TF motifs variable between alpha or beta cell states, we performed two-sided Student’s T-test on motif z-scores between cells labeled as alpha 1 (GCG^high^) and alpha 2 (GCG^low^) cells or beta 1 (INS^high^) and beta 2 (INS^low^). We adjusted raw p-values with the Benjamini-Hochberg procedure to obtain FDR. Motifs with FDR less than 0.05 and absolute difference (Δ) in z-score (between GCG^high^/GCG^low^ alpha or INS^high^/INS^low^ beta) greater than 0.5 were defined as differential motifs (N=46 for beta cells, N=109 for alpha cells and N=111 motifs combined). For these 111 motifs that were variable between alpha or beta cell states, we summarized the mean z-scores over GCG^high^, GCG^low^, INS^high^ and INS^low^ cells and plotted the normalized value. In order to check how motif usage changed along the trajectories, we smoothed motif z-scores along the trajectory for alpha and beta cells separately at step=0.05, using the shrinkage version of cubic regression spline (‘gam’ function from the R package ‘mgcv’ (v1.8.28) with parameter bs=’cs’). We then smoothed motif enrichment profiles and normalized values for visualization. We identified specific TFs likely driving enrichments for a given motif through high Spearman correlation (σ>.9) between motif enrichment and promoter accessibility across the trajectory.

To analyze differential promoter accessibility between alpha and beta cell states, we first calculated the binary promoter by cell matrix containing information about read overlap per cell in a promoter peak. Based on this matrix and cell cluster labels, we performed two-sided Fisher’s exact tests between hormone-high and hormone-low states of alpha, beta, and delta cells for each promoter against the null hypothesis that the promoter had similar accessibility across states. We used Bonferroni adjusted p-values (adjusted p-value<0.01) for alpha and beta cells with the sign of the log_2_ transformed odds ratio to identify genes whose promoter had either increased or decreased accessibility across states. Differentially-accessible promoters were further input into Enrichr^103^ (v.1.0) to perform GO term enrichment analysis on biological processes terms (2018 version). To identify more specific processes, we filtered for gene ontology terms that contained less than 150 total genes.

To plot the profile of each promoter across pseudo-state, we first binned alpha cells or beta cells to 100 bins along the state trajectory. For each bin, we calculated the fraction of cells had a peak in the promoter region for each promoter. Then we smoothed these 100 fractions using the ‘loess’ function from R. The smoothed data were then normalized and clustered using k-medoids clustering, with k determined by optimum average silhouette width using the ‘pamk’ function from the R ‘fpc’ package (v.2.1.11.1). Genes attributed to the promoters in each cluster were then used to perform GO term enrichment analysis.

In order to compare with previous published data, we collected gene lists from Xin et al.^23^. We obtained four gene lists for Beta 1-4 subpopulations (Supplementary Table S3 in Xin et al.). For each gene list, we performed gene set enrichment analysis^104^ using significantly differential promoters (from Figure 2a) as the gene lists to assess whether alpha and beta cell states showed concordant differences (i.e. differential promoters for GCG^low^ and GCG^high^ alpha to compare beta cell states and vice versa for alpha cells).

### Ordering alpha and beta cells along a trajectory and finding dynamic peaks

We used Cicero^6^ (v.1.1.5) to order all alpha and beta cells along separate trajectories. We started with a sparse binary matrix encoding overlap between the superset of islet regulatory peaks and cells. We extracted all cells belonging to alpha cell sub-clusters and filtered out peaks that were not present in alpha cells. We used the aggregate_nearby_peaks function from Cicero to find peaks within 10 kb and merging their counts to make an aggregate matrix. We then chose peaks to define progress with the aggregated matrix by using the differentialGeneTest function from monocle2^47^ to search for peaks that were differentially accessible between the GCG^high^ and GCG^low^ states (FDR<.1), while modeling total peaks in each cell as a covariate. We then used DDRTree to reduce dimensions and ordered cells along the trajectory, setting the root position as the state with the highest glucagon promoter accessibility. We grouped cells into 10 bins based on their trajectory values. Then we repeated the same procedure for beta cells, with the modification of setting the root position by insulin promoter accessibility.

### GWAS enrichment with aggregate peak annotations

We used cell type specific (CTS) LD score regression^69,105^ (v.1.0.0) to calculate enrichment for GWAS traits. We obtained GWAS summary statistics for quantitative traits related to diabetes^13,55–59^, diabetes^10^, and control traits including psychiatric and autoimmune diseases^60–67^. We prepared summary statistics to the standard format for LD score regression. We used peaks from aggregated reads for each cluster as a binary annotation, and the superset of islet regulatory peaks as the background control. For each trait, we then used CTS LD score regression to estimate the enrichment coefficient of each annotation jointly with the background control.

### GWAS enrichment with single cell annotations

We determined genetic enrichment of accessible chromatin profiles in individual cells. We first split the genome into 5 kb windows and removed windows overlapping blacklisted regions from ENCODE. We created a sparse *m* x *n* matrix containing read depth for *m* cells passing read depth thresholds at *n* windows, and extracted highly variable (HV) windows using mean read depths and normalized dispersion. We then retained genetic variants mapping in HV windows with minor allele frequency [MAF]>.05 mapping outside of the major histocompatibility complex region (MHC, defined by chr6:25,000,000-35,000,000 in hg19 coordinates).

As the accessible chromatin profiles from an individual cell are sparse, we used the bagging algorithm in the make_cicero_cds function from Cicero^6^ to aggregate cells into groups of 10. For each aggregate cell group, we created a binary annotation based on mapped reads for cells in the aggregate. We also created baseline annotations consisting of pooled islet cell type accessible chromatin sites and the 53 baseline v1.1 annotations from LD score regression^68^. We then annotated all variants in HV windows with the aggregate cell and baseline annotations. We determined enrichment of HV variant annotations for fasting glucose level^56^, type 2 diabetes^10^, and two control traits, major depressive disorder^66^ and lupus^63^ GWAS data. In order to correct for the confounding effects of linkage disequilibrium (LD), we performed LD pruning of GWAS data for each trait by first sorting variants based on p-value and iteratively removing variants in LD (r^2^>.5, 1000 Genomes European subset) with a more significant variant. To then perform enrichment tests on pruned GWAS data we used a previously described method polyTest^106^ to jointly model the annotation for each aggregated cell group with the baseline pooled site and 53 annotations from LD score baseline v1.1. We then calculated a z-score for each aggregate cell based on the effects and standard error from the resulting model. As the grouping method for Cicero uses bootstrap aggregation, a given cell was potentially assigned to multiple aggregates. We therefore calculated an enrichment z-score for each individual cell by averaging enrichment z-scores for each cell across its respective aggregates.

To identify TFs correlated with trait enrichments, we calculated the Spearman correlation coefficient between fasting glucose or type 2 diabetes single cell GWAS enrichment z-scores and chromVAR motif enrichment z-scores using data from all cells or within beta cells. Within each trait, we used Bonferroni correction to adjust correlation p-values for multiple tests.

### Mapping allelic imbalance within clusters

Genomic DNA for genotyping was extracted either from spare islet nuclei (donors 1 and 2), or acinar cells (donor 3). Genomic DNA was extracted using the DNeasy Blood & Tissue Kits (Qiagen) according to manufacturer’s protocol for purification of total DNA from animal blood or cells. Extracted genomic DNA was used for genotyping on the Illumina Infinium Omni2.5-8 v1.4 genotyping array. For genotypes that passed quality filters (non-missing, MAF>.01 in European or African populations in 1KGP), we then imputed genotypes into the HRC reference panel r1.1^107^ using the Michigan Imputation Server^108^. Post-imputation, we removed genotypes with low imputation quality (R^2^<.3). As an additional filter to remove potential false positive heterozygote genotype calls, we removed variants that had greater than 20 read coverage without reads for both alleles. Using cluster assignments for each cell, we split mapped reads for each donor into cluster-specific reads. For cluster-specific reads, we used the WASP pipeline^109^ (v.0.3.0) to correct for reference mapping bias at heterozygous variants. We then used a two-sided binomial test to assess imbalance at heterozygous variants, assuming a null hypothesis where both alleles were equally likely to be observed. For each variant, we then calculated combined imbalance z-scores across donors using Stouffer’s z-score method and used sequencing depth to weight statistics from each sample.

### Predicting genetic variant effects on chromatin accessibility

We used deltaSVM^70^ to predict the effects of non-coding variants on chromatin accessibility in each cell type and cell state. We obtained sequences underlying promoter-distal (>+/-500 bp from GENCODEv19 transcript TSS for protein coding and long non-coding RNA genes) peaks for each cluster, used ‘genNullSeqs’ to generate background sequences, and then trained a model for each cluster with ‘gkmtrain’ with default settings. For all possible combinations of 11mers, we then used ‘gkmpredict’ to predict the effects of 11mers based on the trained model for the cluster. For each SNP in the HRC reference panel r1.1^107^ overlapping an islet cell type accessible chromatin site, we created 19 bp sequences around each allele (9 bp flanking either side of the variant base). We then used the ‘deltasvm.pl’ script to calculate deltaSVM scores for differential chromatin accessibility between variant alleles. We built a null distribution by randomly permuting the effects of 11mers and re-calculating deltaSVM scores and using the parameters of this null distribution, we calculated z-scores for each variant. From variant z-scores we calculated p-values and then q-values and considered variants significant at FDR<.1.

For variants with predicted effects on chromatin accessibility in alpha or beta cells, we categorized them based on their effects across cell type and states. Variants with significant effects in both alpha cell states but neither beta cell state were classified as “alpha” (n=10,564) and vice versa for “beta” (n=12,833). Variants with significant effects in GCG^high^ alpha and INS^high^ beta states but not GCG^low^ alpha and INS^low^ beta states were classified as “hormone-high” (n=15,769), and vice versa for “hormone-low” (n=12,471). Variants with significant effects in all four alpha and beta cell states were classified as “shared” (n=31,331). We also determined the concordance in the direction of effect for variants across alpha and beta cell states. For the set of variants with significant effects in each state, we calculated the fraction of variants where the allele with the higher predicted effect had a higher predicted effect in other states. We determined significance using a two-sided binomial test assuming an expected fraction of 50%. We assessed enrichment of predicted effect variants in alpha or beta cell states for islet caQTLs^72^ compared to any islet caQTL in alpha or beta cell sites using two-sided Fisher’s exact tests. We then stratified variants with predicted effects by category (hormone-high, hormone-low, alpha, or beta) and assessed enrichment of caQTLs with predicted effects within each category with two-sided Fisher’s exact tests.

### Luciferase gene reporter assays

We selected fine-mapped T2D risk variants with deltaSVM predictions (rs7482891, rs34584161, rs17712208, rs78840640, rs4679370) to test for allelic differences in enhancer activity in MIN6 beta cells using luciferase reporter assays. We cloned sequences containing reference alleles in the forward orientation upstream of the minimal promoter of firefly luciferase vector pGL4.23 (Promega) using KpnI and SacI restriction sites. For rs7482891 (*TH*) and rs34584161 (*RNF6*), we cloned alternative alleles in the forward direction in the same manner as the reference alleles. For rs17712208 (*PROX-AS1*), rs78840640 (*IGF2BP3*), and rs4679370 (*SLC12A8*), we introduced the alternative alleles via site-directed mutagenesis (SDM) using the NEB Q5 Site-Directed Mutagenesis kit (New England Biolabs). We designed primers using NEBaseChanger (v.1.2.8), and we used 10ng of the reporter plasmid containing the reference allele as a template in site-directed mutagenesis using Q5 Hot Start High-Fidelity master mix (New England Biolabs). 4uL of the SMD PCR product was treated with KLD mix (New England Biolabs) and transformed into DH5a *E. coli*. We miniprepped plasmids using the Qiaprep Spin Mini kit (Qiagen) and verified plasmid sequences through Sanger sequencing with the RV3 primer.

SDM primers:

rs17712208 (*PROX-AS1*)
SDM primer (left): GGAGCTATGGaTAATTATTGACTG
SDM primer (right): ATTAACGATCCAGTCAGC

rs78840640 (*IGF2BP3*)
SDM primer (left): ATCAGATTTGgTGAGAAAGAAGAAC
SDM primer (right): GCCCATCAATTCTGAGCATG

rs4679370 (*SLC12A8*)
SDM primer (left): ATCAGTAAGCcCCTAAAGCCTG
SDM primer (right): TAACTTGAGGCAATGGTG

Construct primers:

rs7482891 (*TH*)
construct primer (left): AGAGGTCTGAGGAGCCCTTG
construct primer (right): TAGACCCTGCAGAGCCACAG

rs34584161 (*RNF6*)
construct primer (left): AAGCTGACAGACAGAGGGTCA
construct primer (right): GGGCTTCATAAACATCAGCA

rs17712208 (*PROX-AS1*)
construct primer (left): AAGCCCACCTTCGTAAACAT
construct primer (right): TGAAGTAGCTCCCAGTGAAGG

rs78840640 (*IGF2BP3*)
construct primer (left): CACAATGAAGCCATGTCCTTT
construct primer (right): TCAGCTTTCTATTTTGGGGAAA

rs4679370 (*SLC12A8*)
construct primer (left): TCAATGTCTACCTCAAAATTCTTTGT
construct primer (right): CACTGCAGCCTTAAACTCCTG

We seeded MIN6 cells into 6 (or 12)-well trays at 1 million cells per well. At 80% confluency, we co-transfected cells with 500ng of the experimental firefly luciferase vector pGL4.23 containing the alternative allele, reference allele, or an empty vector and 50ng of the vector pRL-SV40 (Promega) using the Lipofectamine 3000 reagent (Thermo Fisher). We performed all transfections were performed in triplicate. Six hours after transfection, we replaced MIN6 growth media consisting of modified DMEM containing 1.5g/L sodium bicarbonate supplemented with 4% heat inactivated FBS, gentamicin, and 50uM beta-mercaptoethanol. We lysed cells 48 hours after transfection and assayed for Firefly and Renilla luciferase activities using the Dual-Luciferase Reporter system (Promega). We normalized Firefly activity to Renilla activity and compared it to the empty vector, and normalized results were expressed as fold change compared to empty vector control per allele. We used a two-sided Student’s T-test to compare the luciferase activity between the two alleles.

### TF motif enrichment within predicted effect variant categories

For each cell- or state-resolved category (hormone-high, hormone-low, alpha, beta) of variants with predicted effects, we extracted 29 bp sequences (+/-14 bp around each SNP) corresponding to the higher or lower predicted effect allele. Here, we reasoned that extracting sequences for a larger window around SNPs would alleviate bias for the analysis against motifs with longer PWMs. We then used AME from the MEME suite^110^ (v.4.12.0) to predict motif enrichment, using position weight matrices from the latest non-redundant motif library JASPAR 2018^38^. We used sequences from the higher effect allele as the test set and sequences from the lower effect allele as the background set. Because motif for TFs within the same structural family can potentially show similar enrichment, we used the TFClass database (http://tfclass.bioinf.med.uni-goettingen.de/) to group motifs by TF family. To determine which TF was most likely driving the enrichment, we used min-max normalized promoter accessibility within TF family members with a promoter peak in alpha or beta cells and highlighted corresponding cell type patterns of promoter accessibility.

### Enrichment of predicted variants for lower frequency variants

We obtained genome-wide summary statistics of T2D from the DIAMANTE consortium^10^. We estimated LD patterns for variants with MAF<.05 using HRC imputed genotype data from samples in the UK Biobank (UKB, March 2018 release). We randomly selected 10,000 non-related UKB samples of European ancestry and calculated LD between lower frequency variants using PLINK (v.1.90b6.7). We then LD-pruned variants with MAF<.05 in DIAMANTE T2D data by first sorting variants based on their p-values and then removed variants in r^2^>.5 with a more significant variant. Using the LD-pruned results, we then determined enrichment of variants with predicted effects on endocrine cell types. We created sets of variants that had significant effects (FDR<.1) in any endocrine cell type, as well as variants with FDR<.1 for each cell type. For alpha, beta and delta cells, we considered variants with effects in either cell state. We then created a background set of variants as those without significant effects in any endocrine cell type (all FDR>.1). We set a series of p-value thresholds (5×10^−8^, 1×10^−7^, 1×10^−6^, 1×10^−5^, 1×10^−4^, 1×10^−3^), and at each threshold determined the fraction of variants in each category as well as background variants reaching that p-value threshold to calculate a fold-enrichment based on these fractions compared to background. We determined significance of the enrichments by using a two-sided binomial test of the counts for each category using the background fraction as the expected count.

### Chromatin co-accessibility with Cicero

We used Cicero^6^ to calculate peak-peak co-accessibility scores for alpha and beta cells. Like the trajectory analysis, we started with a sparse binary matrix encoding overlap between the superset of islet regulatory peaks and cells. We extracted all cells belonging to alpha cell sub-clusters and filtered out peaks that were not present in alpha cells. We then used the make_cicero_cds function to aggregate cells based on the 50 nearest neighbors from the UMAP reduced dimensions. We then used Cicero to calculate co-accessibility scores using a window size of 1 Mb and a distance constraint of 500 kb, leaving other parameters at the default setting. We then repeated the same procedure for beta cells. We used two-sided Fisher’s exact tests to assess whether distal co-accessible sites had higher accessibility in either hormone-high or -low states, and defined significance at FDR<.1. To compare promoter-distal co-accessibility links that had higher accessibility in the same direction (either both hormone-high or hormone-low), we used differential promoters between states (from the previous analysis in Figure 2).

### Enrichment of islet Hi-C and pcHi-C loops in co-accessible peaks

We obtained sets of merged Hi-C loops^27^ and high-confidence promoter capture Hi-C (pcHi-C) loops^76^ from public datasets. For Hi-C loops, we used anchors directly from the loops. For pcHi-C loops, we used a 5 kb window centered on the interaction point as the anchor. To compare alpha and beta cell co-accessibility with Hi-C, we then used direct overlap of alpha or beta cell peaks with anchors. For different binned thresholds of co-accessibility in .05 increments, we then calculated distance-matched odds ratios for co-accessible peaks containing Hi-C loops versus non-co-accessible peaks (co-accessibility<0). We then used two-sided Fisher’s exact tests to assess significance. We repeated the procedure for high confidence pcHi-C loops for both cell types.

### Annotating fine-mapped diabetes risk variants

We annotated risk signals in compiled fine-mapping data for type 2 diabetes from the DIAMANTE consortium and Biobank Japan studies. For the Biobank Japan T2D GWAS, we constructed LD-based 99% genetic credible sets for main signals at 22 novel loci that were distinct from the DIAMANTE study. We used the East Asian subset of the 1000 Genomes Project to define credible set variants by taking all variants in at least low LD (r^2^>.1) with the index variant in a 5 Mb window. We used effect size and standard error estimates to calculate Bayes factors for each variant. For each signal, we then calculated the posterior probability causal probability (PPA) that each variant drives the association by dividing its Bayes factor by the sum of Bayes factors for all variants in the signal’s credible set. We then sorted each signal by descending PPA and retained variants that added up to a cumulative probability of .99 to derive 99% credible sets.

For each signal, we identified candidate casual variants that were both in the 99% credible set and had a posterior causal probability greater than .01. We intersected these candidate variants with accessible chromatin sites for each islet cell type and cell state, and then identified variants with predicted effects on the overlapping cell types/states. We finally annotated variants based on overlap with sites co-accessible to gene promoters. For target genes linked to diabetes risk variants we determined enriched gene sets using GSEA.

### Analysis of *INS* promoter 4C data

We downloaded and re-analyzed published 4C data of the *INS* promoter for the beta cell line EndoC-βH1^77^ with 4C-ker^111^. We first created a reduced genome using 25 bp flanking sequences of BglII cutting sites. For each of the 3 replicates, we then aligned reads to this reduced genome using bowtie2^112^ (v.2.2.9) with the parameter “-N 0 −5 20”. We then extracted counts for each fragment from the SAM file after removing self-ligated and undigested fragments, and we used the bedGraph files as input to the R.4Cker package. We generated normalized counts and called high interaction regions using the ‘nearBaitAnalysis’ function with the parameter ‘k=10’.

### CRISPR/Cas9-mediated enhancer deletion

H1 hESCs (WA01; purchased from WiCell; NIH registration number: 0043) were seeded onto Matrigel®-coated six-well plates at a density of 50,000 cells/cm^2^ and maintained in mTeSR1 media (StemCell Technologies) for 3-4 days with media changed daily. hESC research was approved by the University of California, San Diego, Institutional Review Board and Embryonic Stem Cell Research Oversight Committee.

To generate clonal homozygous *KCNQ1* enhancer deletion hESC lines, two sgRNAs targeting the enhancer were designed and cloned into Px333-GFP, a modified version of Px333 (#64073, Addgene). The plasmid was transfected into H1 hESCs with XtremeGene 9 (Roche). 24 hours later, 5000 GFP^+^ cells were sorted into a well of six-well plate. Individual colonies that emerged within 5-7 days after transfection were subsequently transferred manually into 48-well plates for expansion, genomic DNA extraction, PCR genotyping, and Sanger sequencing. sgRNA oligos and genotyping primers are listed below. For control clones, we transfected the Px333-GFP plasmid into H1 hESCs and subjected the cells to the same workflow as H1 hESCs transfected with sgRNAs.

sgRNA oligos:

KCNQ1_sgRNA1-s: ACTGTCGGGCCCATCTGCCA
KCNQ1_sgRNA1-as: TGGTTGGATCTGTTGCGGGG

Genotyping primers:

Span-F: AGTGGGGCCATGAACAATAA
Span-R: GCCTGAGTTTCCGTGACTGT

### Pancreatic differentiation of enhancer-deleted hESCs clones

hESCs were differentiated in a suspension-based format using rotational culture with some modifications to a published protocol^83^. Undifferentiated hESCs were aggregated by preparing a single cell suspension in mTeSR media (STEMCELL Technologies) at 1×10^6^ cells/mL and overnight culture in six-well ultra-low attachment plates (Costar) with 5.5ml per well on an orbital rotator (Innova2000, New Brunswick Scientific) at 100 rpm. The following day, undifferentiated aggregates were washed in DMEM/F12 (VWR) and differentiated using a multistep protocol with daily media changes and continued orbital rotation at either 100 rpm or at 108 rpm from days 8 to 28. In addition to 1% GlutaMAX™ (Gibco) and 15 mM (days 0-10) or 20 mM (days 11-28) glucose, MCDB 131 media (Life Technologies) was supplemented with 0.5% (days 0-5) or 2% (days 6-14) fatty acid-free BSA (Proliant), 1.5 g/L (days 0-5 and days 11-28) or 2.5 g/L (days 6-10) NaHCO_3_ (Sigma-Aldrich), and 0.25 mM (days 3-10) ascorbic acid (Sigma-Aldrich).

Human Activin A, mouse Wnt3a, and human KGF were purchased from R&D Systems. Other media components included ascorbic acid (Sigma-Aldrich), Insulin-Transferrin-Selenium-Ethanolamine (ITS-X; Thermo Fisher Scientific), ZnSO_4_ (Sigma-Aldrich), heparin (Sigma-Aldrich), retinoic acid (RA) (Sigma-Aldrich), SANT-1 (Sigma-Aldrich), 3,3′,5-Triiodo-L-thyronine (T3) (Sigma-Aldrich), the protein kinase C activator TPB (EMD Chemicals), the BMP type 1 receptor inhibitor LDN-193189 (Stemgent), the TGFβ type 1 activin like kinase receptor ALK5 inhibitor, ALK5 inhibitor II (Enzo Life Sciences), N-Acetyl-L-cysteine (Sigma), R428 (SelleckChem), Trolox (EMD Millipore), γ-secretase inhibitor XX (Calbiochem).

Day 0: MCDB 131, 100ng/mL Activin, 25ng/mL mouse Wnt3a

Day 1 – Day 2: MCDB 131, 100ng/mL Activin A

Day 3 – Day 5: MCDB 131, 50ng/mL KGF

Day 6 – Day 7: MCDB 131, 50ng/mL KGF, 0.25 µM SANT-1, 1 µM RA 100 nM LDN-193189, 200 nM TPB, 0.5% ITS-X

Day 8 – Day 10: MCDB 131, 2ng/mL KGF, 0.25 µM SANT-1, 0.1 µM RA, 200 nM LDN-193189, 100 nM TPB, 0.5% ITS-X

Day 11 – Day 13: MCDB 131, 0.25 µM SANT-1, 0.05 µM RA, 100 nM LDN-193189, 1 µM T3, 10 µM ALK5i II, 10 µM ZnSO_4_, 10 µg/mL heparin, 0.5% ITS-X

Day 14 – Day 21: MCDB 131, 100 nM LDN-193189, 1 µM T3, 10 µM ALK5i II, 10 µM ZnSO_4_, 10 µg/mL heparin, 100nM γ-secretase inhibitor XX, 0.5% ITS-X

Day 21 – Day 28: MCDB 131, 100 nM LDN-193189, 1 µM T3, 10 µM ALK5i II, 10 µM ZnSO_4_, 10 µg/mL heparin, 1mM N-Acetyl-L-cysteine, 10µM Trolox, 2µM R428, 0.5% ITS-X

### Characterization of hESC-derived cultures at beta cell stage (day 28)

#### Flow cytometry analysis

hESC-derived cell aggregates were dissociated into a single-cell suspension with Accutase™ (Innovative Cell Technologies) at 37 °C for 5 min. Accutase™ was quenched with FACS buffer (0.2% (w/v) BSA in PBS). Cells were then pelleted, fixed, and permeabilized with Cytofix/Cytoperm Fixation/Permeabilization Solution (BD Biosciences) for 20 min at 4 °C, and washed twice in BD Perm/Wash™ Buffer. We incubated cells with AF647-conjugated mouse anti-Nkx6.1 (BD Biosciences) and PE-conjugated rabbit anti-INS (Cell Signaling Technology) antibody in 50 µl BD Perm/Wash™ Buffer for 1 hour at 4 °C. Following three washes in BD Perm/Wash™ Buffer, cells were analyzed on a FACSCanto II (BD Biosciences) cytometer.

#### Immunofluorescence staining and quantification of immunofluorescence signal

hESC-derived cell aggregates were washed twice with PBS and then fixed with 4% paraformaldehyde in PBS for 30 min at room temperature. Following three washes in PBS, aggregates were incubated in 30% sucrose at 4 °C overnight, frozen in Optimal Cutting Temperature Compound (Sakura Finetek USA), and sectioned at 10 µm with a CM3050S cryostat (Leica). Sections were washed with PBS, permeabilized, and blocked with Permeabilization/Blocking Buffer for 1 h at room temperature. Primary and secondary antibodies were diluted in Permeabilization/Blocking Buffer. We incubated sections overnight at 4°C with primary antibodies, and then secondary antibodies for 30 min at room temperature. The following primary antibodies were used: mouse anti-NKX6-1 (LifeSpan BioSciences, 1:250), guinea pig anti-INS (Dako, 1:1000). Secondary antibodies (1:1000) were Cy3-, Alexafluor488-conjugated antibodies raised in donkey against mouse and guinea pig (Jackson Immuno Research Laboratories). We acquired images on a Zeiss Axio-Observer-Z1 microscope with a Zeiss AxioCam digital camera.

#### mRNA sequencing

For each clone, we collected aggregates from two independent batches of differentiation and lysed them in RLT Buffer. We then extracted total RNA using the RNeasy Micro Kit (QIAGEN) following the manufacturer’s instructions. mRNA libraries were prepared using KAPA mRNA Hyper Prep kit (KAPA) and single-end 75 bp reads were sequenced using HiSeq4000 (Illumina). We used STAR (v2.5.3a) to map reads to the hg19 genome, allowing for up to 10 mismatches. We retained reads aligned uniquely to one genomic location for subsequent analysis. We then created input count files for DESeq2 with htseq-count from the HTSeq python package (v.0.9.0) and tested for differential gene expression using DESeq2 (v1.10.1) with default parameters, using differentiation batch as a technical covariate in our analysis. We considered genes with an FDR<.1 as significantly differentially expressed.

#### Insulin content measurement

We washed hESC-derived cell aggregates with PBS, resuspended in 50μl of 0.1% SDS TE buffer and sonicated for 3 cycles of 30 sec on/ 30 sec off each using a Bioruptor on the high setting. We then immersed the lysate in a solution of 2% HCl and 80% ethanol overnight at 4°C and centrifuged at max speed for 10 min at 4°C. We collected the supernatant and measured insulin content using a human insulin ELISA kit (ALPCO). We resuspended the pellets in 50µl TE buffer and measured DNA content with Nanodrop, and normalized insulin content to DNA content.

## Acknowledgements

This work was supported by NIH R01DK114650 and U01DK105554 (sub-award) to K.G., R01DK068471 and U01DK105541 to M.S., U01DK120429 to K.G. and M.S., and by the UC San Diego School of Medicine to the Center for Epigenomics. Data from the UK Biobank was accessed under application 24058. We thank Ileana Matta for preparation of RNA-seq libraries.

## Conflict of Interest

The authors have no conflict of interest to disclose.

## Author Contributions

K.J.G., D.U.G, and M.Sa. conceived of and supervised the research in the study; K.J.G., D.U.G., M.Sa., J.C., C.Z, and Z.C. wrote the manuscript; J.C. performed analyses of single cell and genetic data; C.Z., M.Sc and J.W. performed hESC experiments; Z.C. performed analyses of single cell data; J.Y.H. performed single cell assays and genotyping; S.H., A.D. and M.O. performed reporter experiments; Y.Q. performed analyses of 4C data; Y.Sui performed analyses of hESC data; Y.Sun and P.K. developed and processed data for the epigenome database; R.F. contributed analyses of single cell data; S.P. contributed to the development of single cell assays.

## Data Availability

Processed data and annotations will be made available at www.t2depigenome.org, and raw sequence data will be deposited in GEO prior to publication. All other source data are either included in the study or available from the corresponding authors upon request.

