## Supplementary Figures for "Single cell chromatin accessibility reveals pancreatic islet cell type- and state-specific regulatory programs of diabetes risk"

### Figure S1

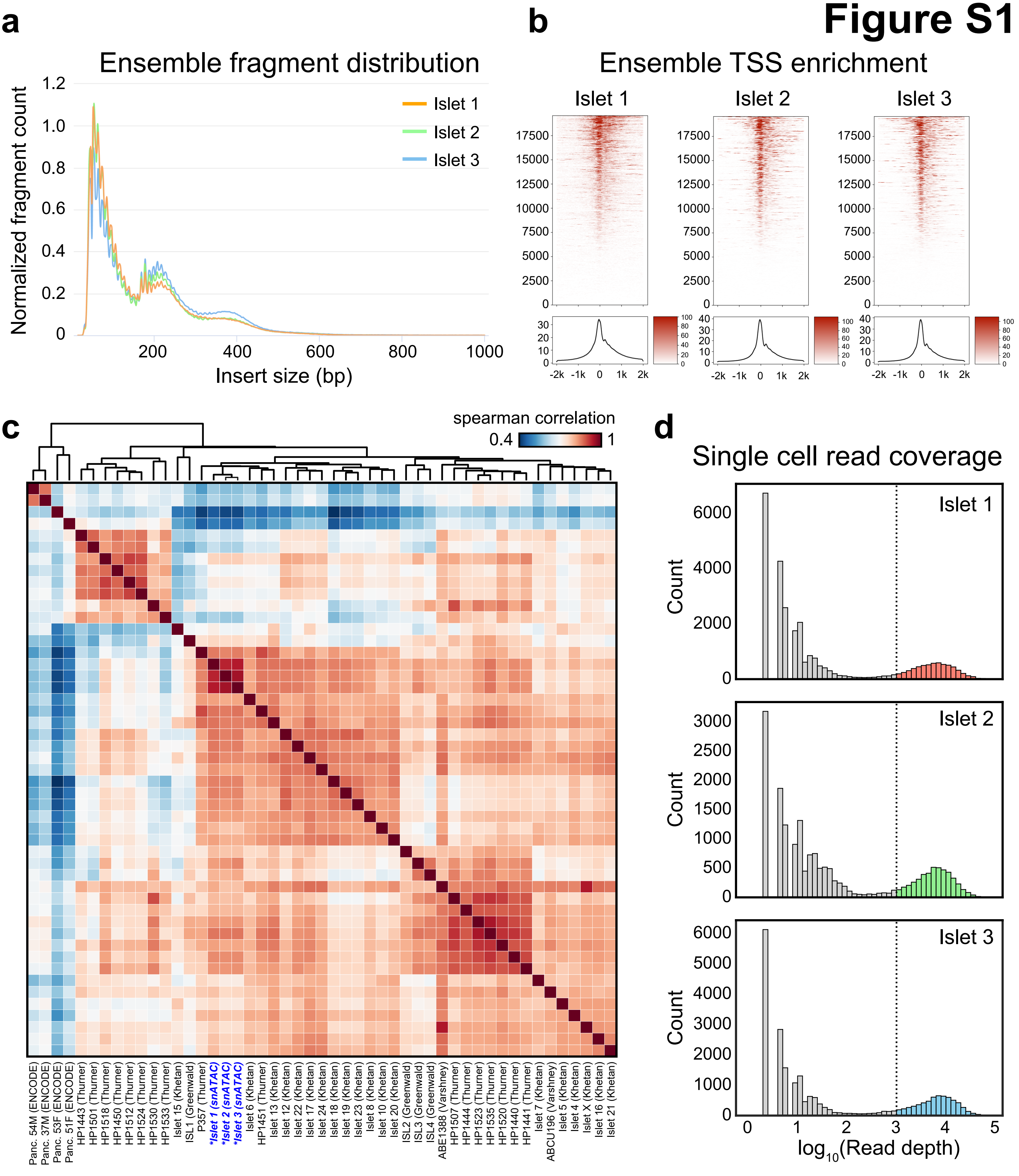


**Supplementary Figure 1. Quality control metrics and aggregate comparison to bulk islet ATAC.** (a) Insert size distribution for aggregate reads from each snATAC-seq experiment. (b) Aggregated read coverage from each snATAC-seq experiment in a +/-2kb window around individual promoters (top) and averaged across all promoters (bottom). (c) Spearman correlation between normalized read coverage within a merged set of peaks from 3 aggregated islet snATAC-seq, 42 bulk islet ATAC-seq, and 4 bulk pancreas ATAC-seq datasets. Names of samples are from the original sources of the data, which are indicated in parentheses. (d) Binned $\log_{10}$ read depth distribution for each experiment.

Figure S2

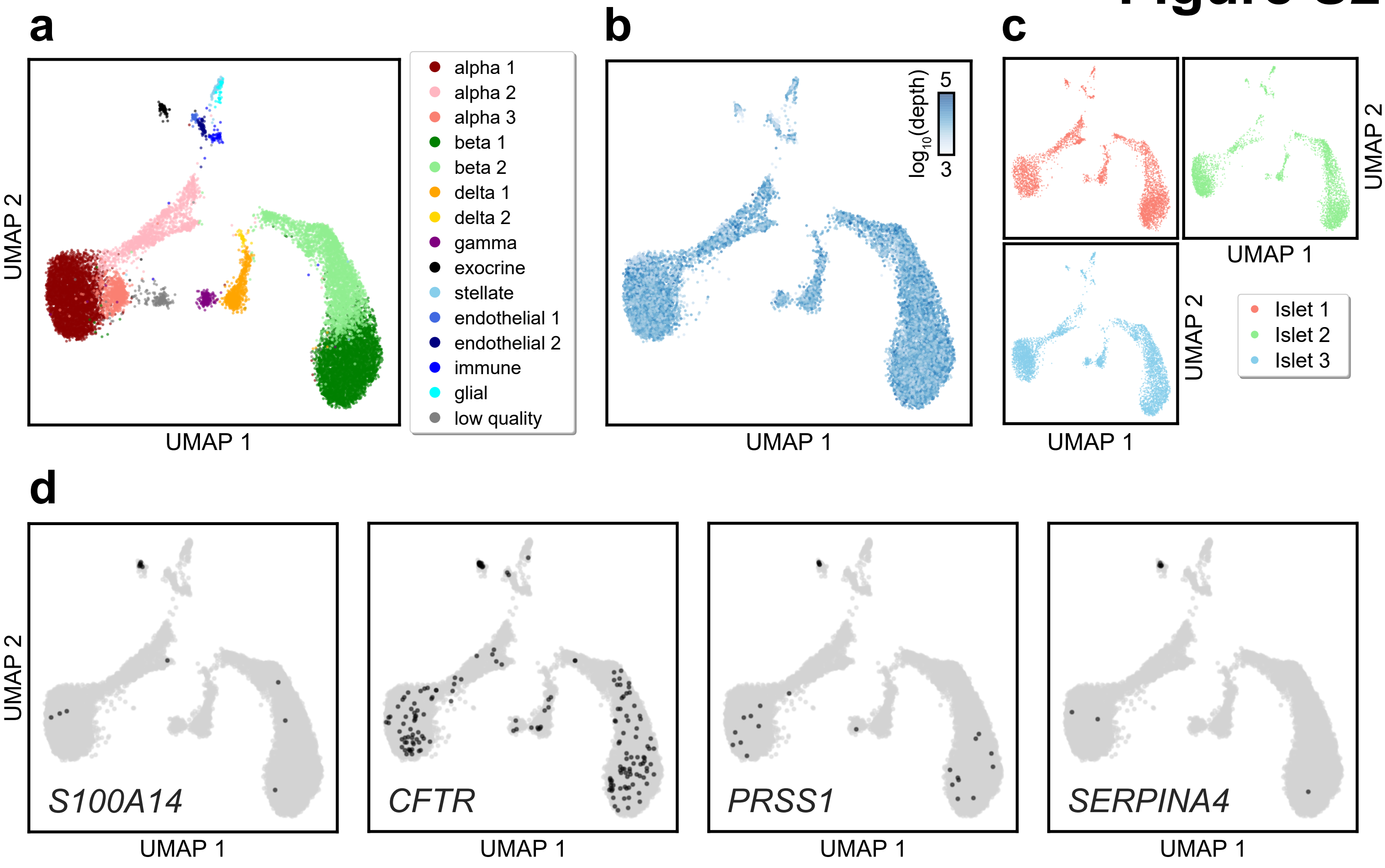


**Supplementary Figure 2. Quality metrics and marker promoters of islet snATAC clusters.** (a) Islet snATAC-seq clusters on UMAP coordinates including alpha 3 and doublets. (b) Log10 normalized total read depth for each cell on UMAP coordinates. (c) Cells colored by donor of origin on UMAP coordinates. (d) Promoter accessibility in a 1kb window around the TSS for pancreatic duct (*S100A14* and *CFTR*) or acinar (*PRSS1*, *SERPINA4*) marker genes for each profiled cell. A cell is colored black if it had promoter accessibility for the marker gene listed in the bottom right corner of each subplot, or otherwise is grey.

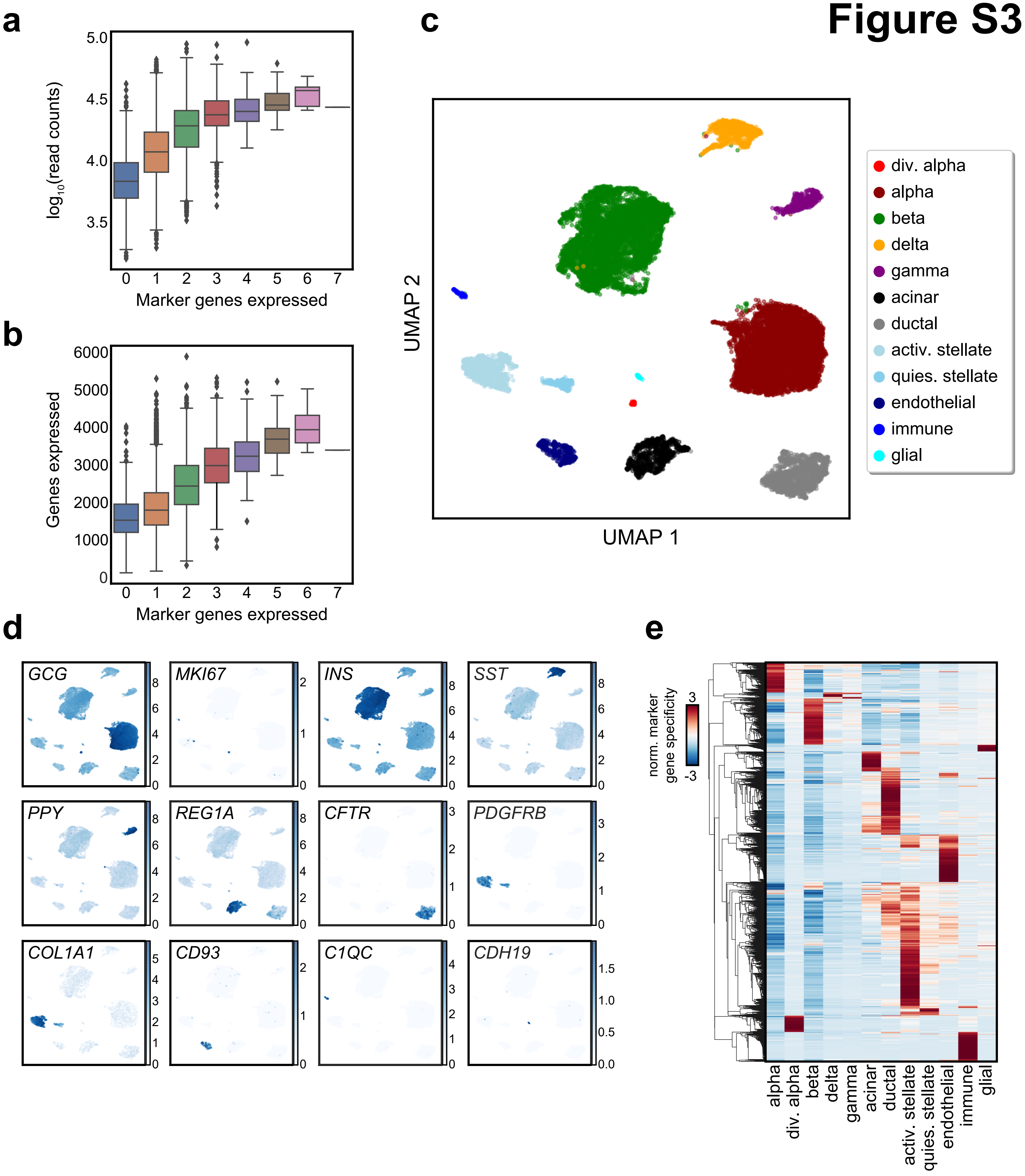


**Supplementary Figure 3. Analysis of islet single cell gene expression data.** (a) $Log_{10}$ transformed read depth or (b) total number of genes expressed compared with number of marker genes expressed per cell from single cell RNA-seq data. Cells expressing more than one marker gene (defined by mixture models) were marked as doublets and filtered out. (c) Clusters of islet cells from single cell RNA-seq data plotted on UMAP coordinates. div. alpha, dividing alpha. quies. stellate, quiescent stellate. activ. stellate, activated stellate. (d) Selected marker gene $log_2$(expression) for each cluster plotted on UMAP coordinates. (e) Row-normalized t-statistics of marker gene specificity showing the most specific genes (t-statistic>20) for each cluster.

### Figure S4

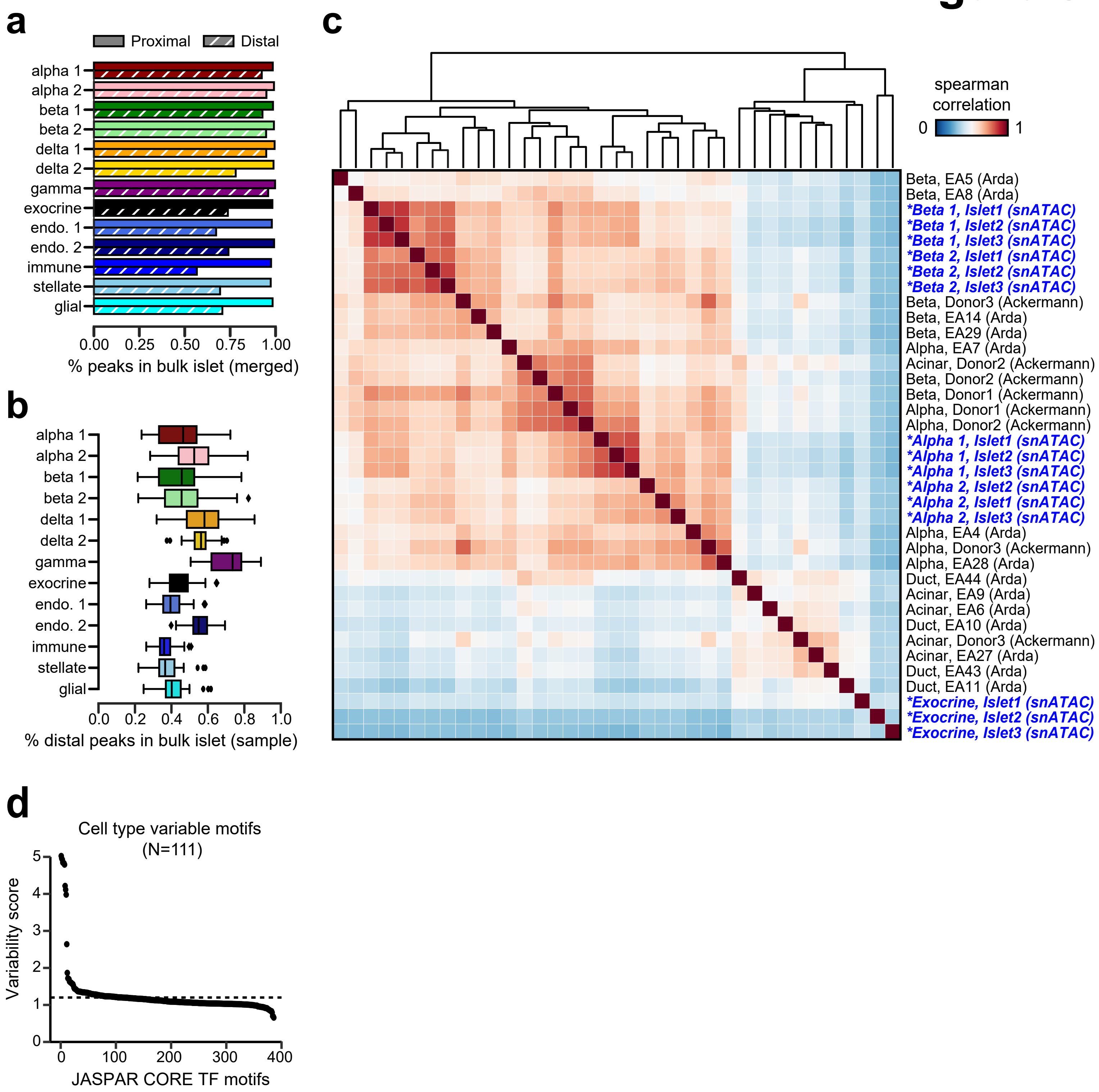


**Supplementary Figure 4. Evaluation of islet snATAC clusters through comparison to ensemble/sorted data and motif enrichment.** (a) Percentage of promoter proximal (solid) or distal (hatched) aggregate peaks for each islet snATAC-seq cluster found in the merged set of bulk islet ATAC-seq peaks. (b) Boxplot showing the distribution of percentage of promoter distal peak overlap for each cluster with each individual bulk islet ATAC-seq sample. (c) Spearman correlation between normalized read coverage within a merged set of peaks from alpha, beta, and exocrine snATAC-seq clusters from 3 snATAC-seq, 6 sorted alpha cell bulk ATAC-seq, 7 sorted beta cell bulk ATAC-seq, 4 sorted ductal cell bulk ATAC-seq, and 5 sorted acinar cell bulk ATAC-seq datasets. Names of samples are from the original sources of the data, which are indicated in parentheses. (d) Selection criteria for 111 JASPAR CORE TF motifs with highly variable chromVAR motif enrichment across islet and pancreatic cell types (related to Fig. 1e).

### Figure S5

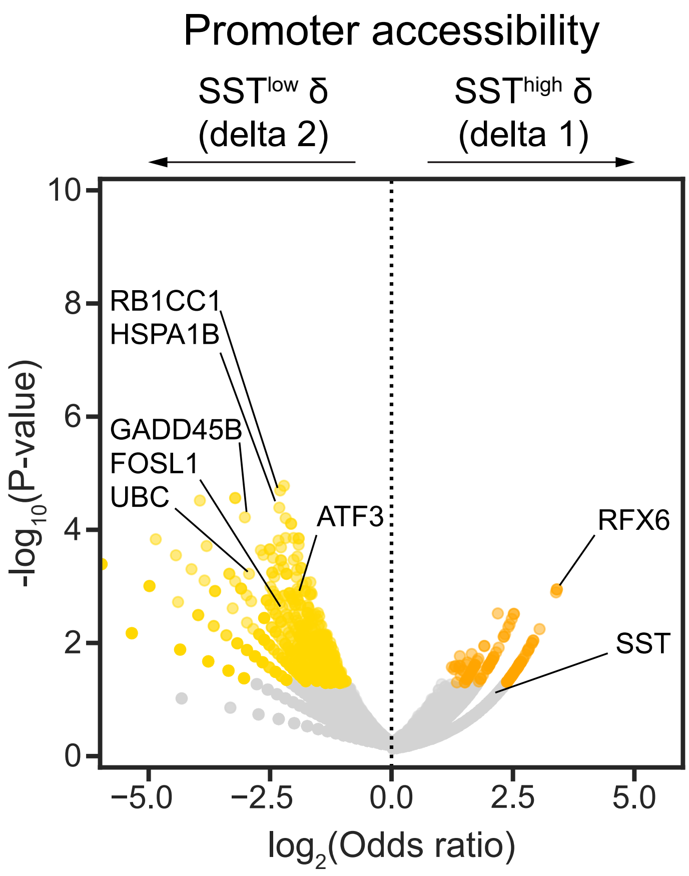


**Supplementary Figure 5. Differentially accessible promoters across states and pseudo-states.** (a) Gene promoters with differential chromatin accessibility at a nominal significance threshold (P<.05). *RFX6* is among genes with increased promoter accessibility in delta 1 ($SST^{high}$ δ), while stress response-related genes such as *FOSL1* and *ATF3* are included among genes with increased promoter accessibility in delta 2 ($SST^{low}$ δ).

### Figure S6

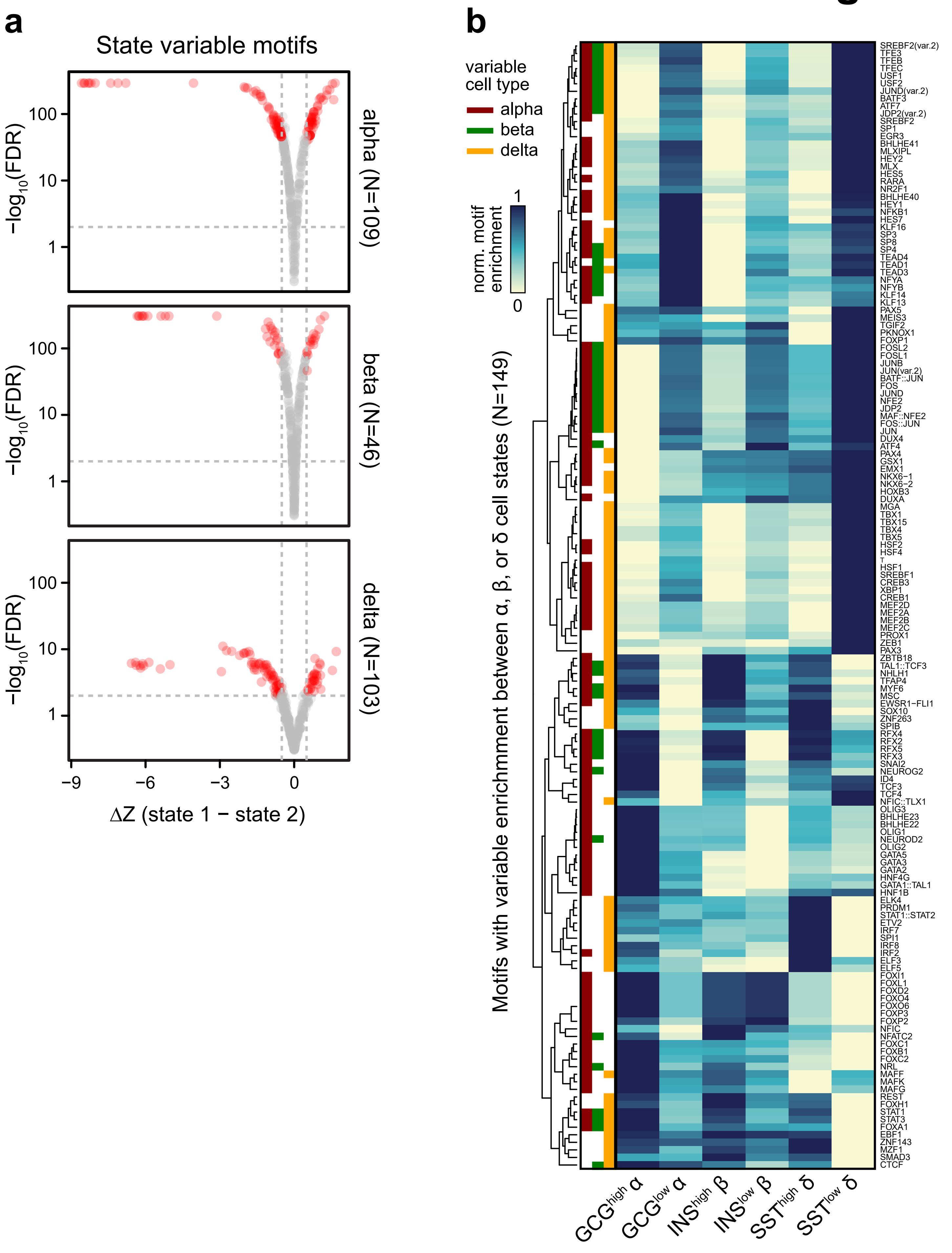

a
State variable motifs
−log10(FDR)
100
10
1
alpha (N=109)
beta (N=46)
delta (N=103)
−9
−6
−3
0
ΔZ (state 1 − state 2)
b
variable cell type
alpha
beta
delta
norm. motif enrichment
1
0
Motifs with variable enrichmment between α, β, or δ cell states (N=149)
GCGhigh α
GCGlow α
INShigh β
INSlow β
SSThigh δ
SSTlow δ
SREBF2(var.2)
TFE3
TFEB
TFEC
USF1
USF2
JUND(var.2)
BATF3
ATF7
JDP2(var.2)
SREBF2
SP1
EGR3
BHLHE41
MLXIPL
HEY2
MLX
HES5
RARA
NR2F1
BHLHE40
HEY1
NFKB1
HES7
KLF16
SP3
SP8
SP4
TEAD4
TEAD1
TEAD3
NFYA
NFYB
KLF14
KLF13
PAX5
MEIS3
TGIF2
PKNOX1
FOXP1
FOSL2
FOSL1
JUNB
JUN(var.2)
BATF::JUN
FOS
JUND
NFE2
JDP2
MAF::NFE2
FOS::JUN
JUN
DUX4
ATF4
PAX4
GSX1
EMX1
NKX6−1
NKX6−2
HOXB3
DUXA
MGA
TBX1
TBX15
TBX4
TBX5
HSF2
HSF4
T
HSF1
SREBF1
CREB3
XBP1
CREB1
MEF2D
MEF2A
MEF2B
MEF2C
PROX1
ZEB1
PAX3
ZBTB18
TAL1::TCF3
NHLH1
TFAP4
MYF6
MSC
EWSR1−FLI1
SOX10
ZNF263
SPIB
RFX4
RFX2
RFX5
RFX3
SNAI2
NEUROG2
ID4
TCF3
TCF4
NFIC::TLX1
OLIG3
BHLHE23
BHLHE22
OLIG1
NEUROD2
OLIG2
GATA5
GATA3
GATA2
HNF4G
GATA1::TAL1
HNF1B
ELK4
PRDM1
STAT1::STAT2
ETV2
IRF7
SPI1
IRF8
IRF2
ELF3
ELF5
FOXI1
FOXL1
FOXD2
FOXO4
FOXO6
FOXP3
FOXP2
NFIC
NFATC2
FOXC1
FOXB1
FOXC2
NRL
MAFF
MAFK
MAFG
REST
FOXH1
STAT1
STAT3
FOXA1
EBF1
ZNF143
MZF1
SMAD3
CTCF

**Supplementary Figure 6. Sequence motifs with differential enrichment across cell states.** (a) Selection of transcription factor motifs with variable enrichment (at $FDR<.05$ and $\Delta Z>.5$) between alpha (109 motifs, top), beta (46 motifs, middle), and delta (103 motifs, bottom) states, many of which were common across islet cell types. The x-axis represents the absolute difference in z-score between $GCG^{high}/GCG^{low}$ α, $INS^{high}/INS^{low}$ β, or $SST^{high}/SST^{low}$ δ. (b) Row-normalized average chromVAR motif enrichments per cell state for 149 motifs with variable enrichment between alpha (α, dark red), beta (β, green), or delta (δ, orange) cell states.

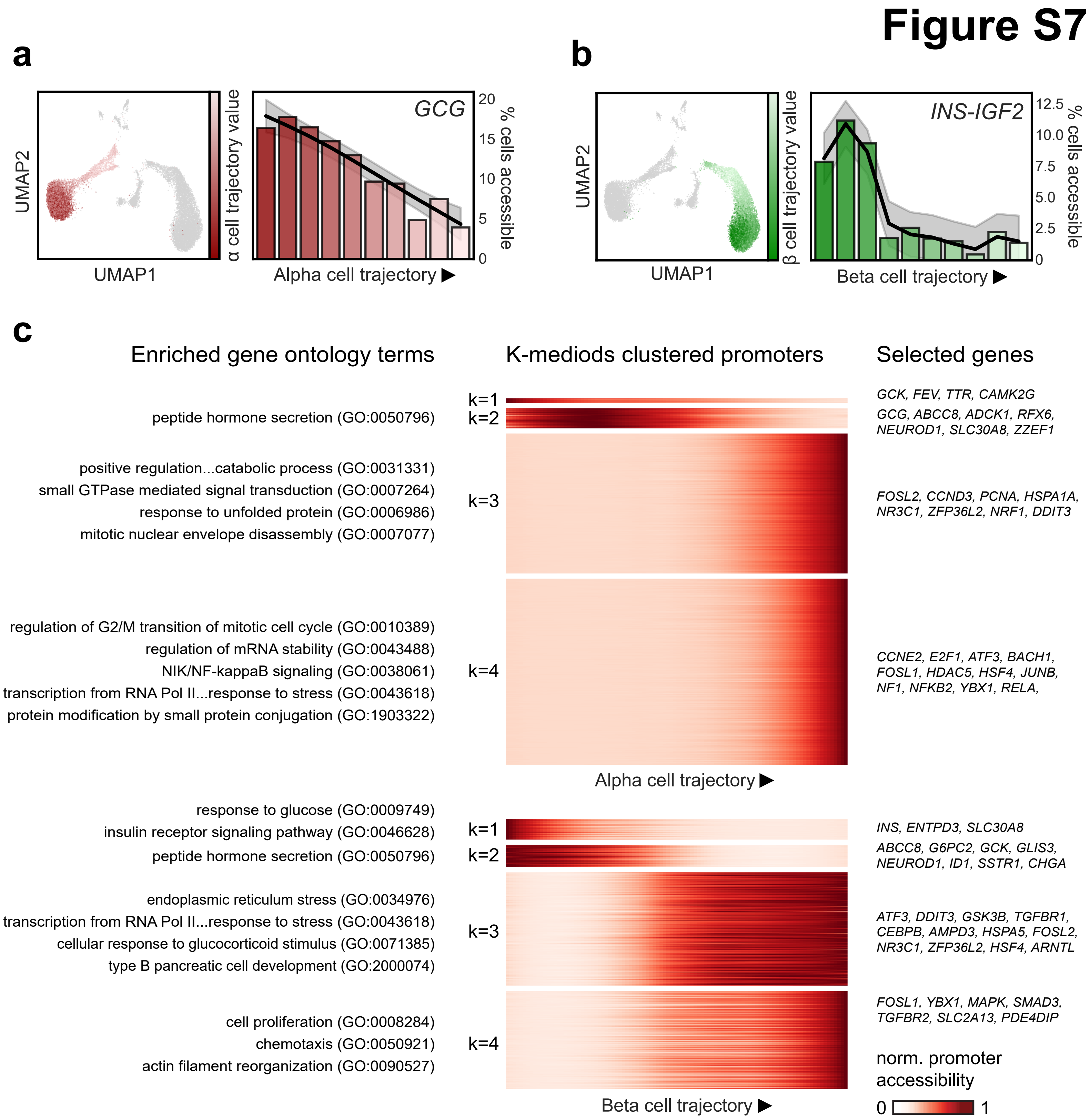


**Supplementary Figure 7. Differentially accessible promoters across pseudo-states.** (a) Pseudo-state (trajectory) values for alpha cells plotted on UMAP coordinates (left) and percentage of cells with *GCG* promoter accessibility decreases across 10 bins along the alpha (α) cell trajectory (right). (b) Pseudo-state (trajectory) values for beta (β) cells plotted on UMAP coordinates (left) and percentage of cells with *INS* promoter accessibility decreases across 10 bins along the beta cell trajectory (right). (c) Heatmaps showing promoters with dynamic accessibility across trajectories for alpha (top) and beta (bottom) cell trajectories. Gene promoters are clustered into 4 groups for each trajectory with k-medoids clustering. Enriched gene ontology for each k-mediod cluster (left) and selected genes present in at least one enriched gene ontology term for the k-medoids cluster.

### Figure S8

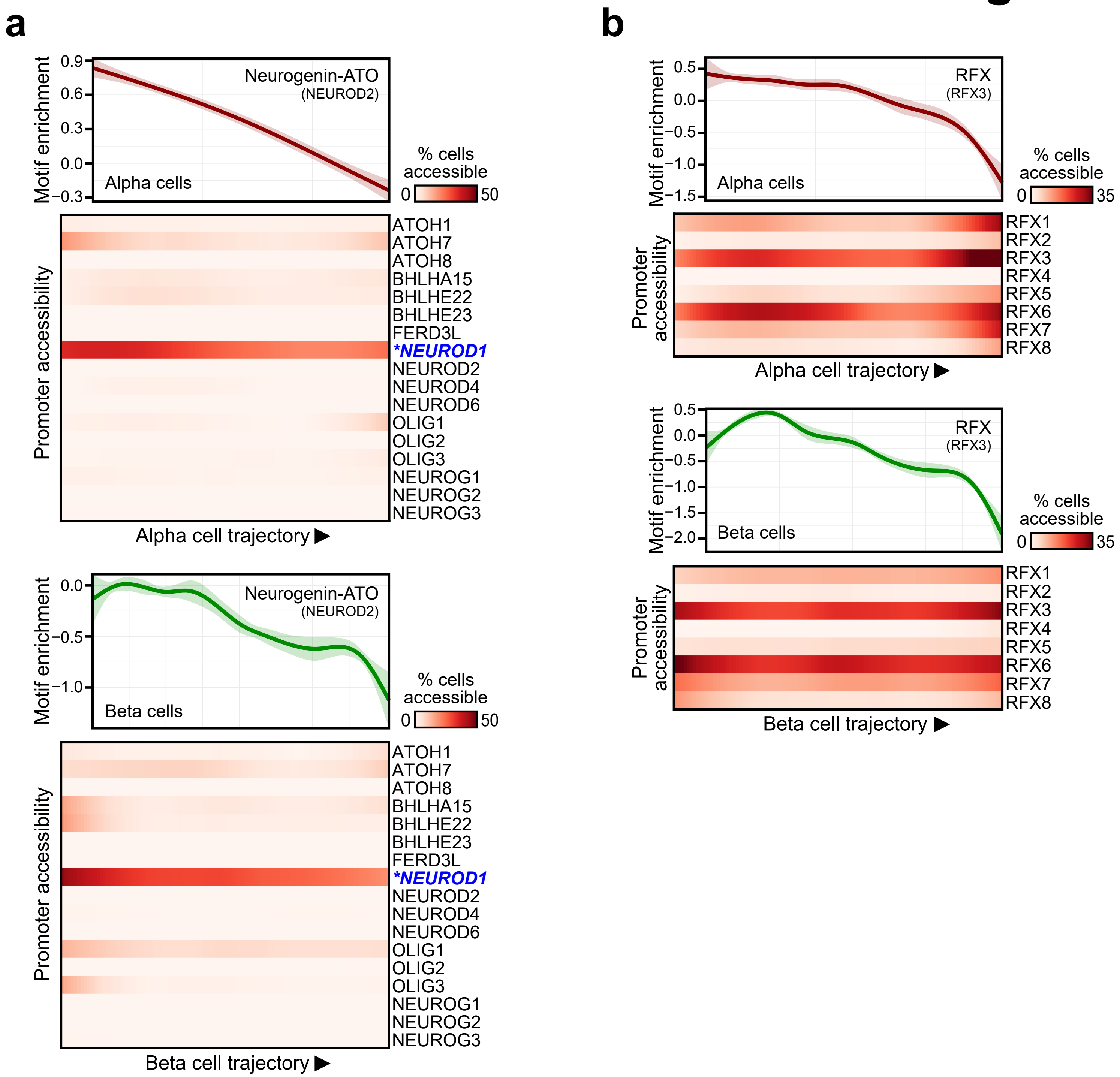


**Supplementary Figure 8. Sequence motifs with dynamic enrichment across alpha and beta cell states.** (a) Motifs in the Neurogenin-ATO subfamily show decreasing enrichment across the alpha (top) and beta (bottom) cell trajectory. NEUROD1 is the only gene in the subfamily with matching patterns of promoter accessibility (Spearman correlation>.9) and is highlighted (in blue and starred). (b) Motifs in the RFX subfamily show decreasing enrichment across the alpha (top) and beta (bottom) cell trajectory.

### Figure S9

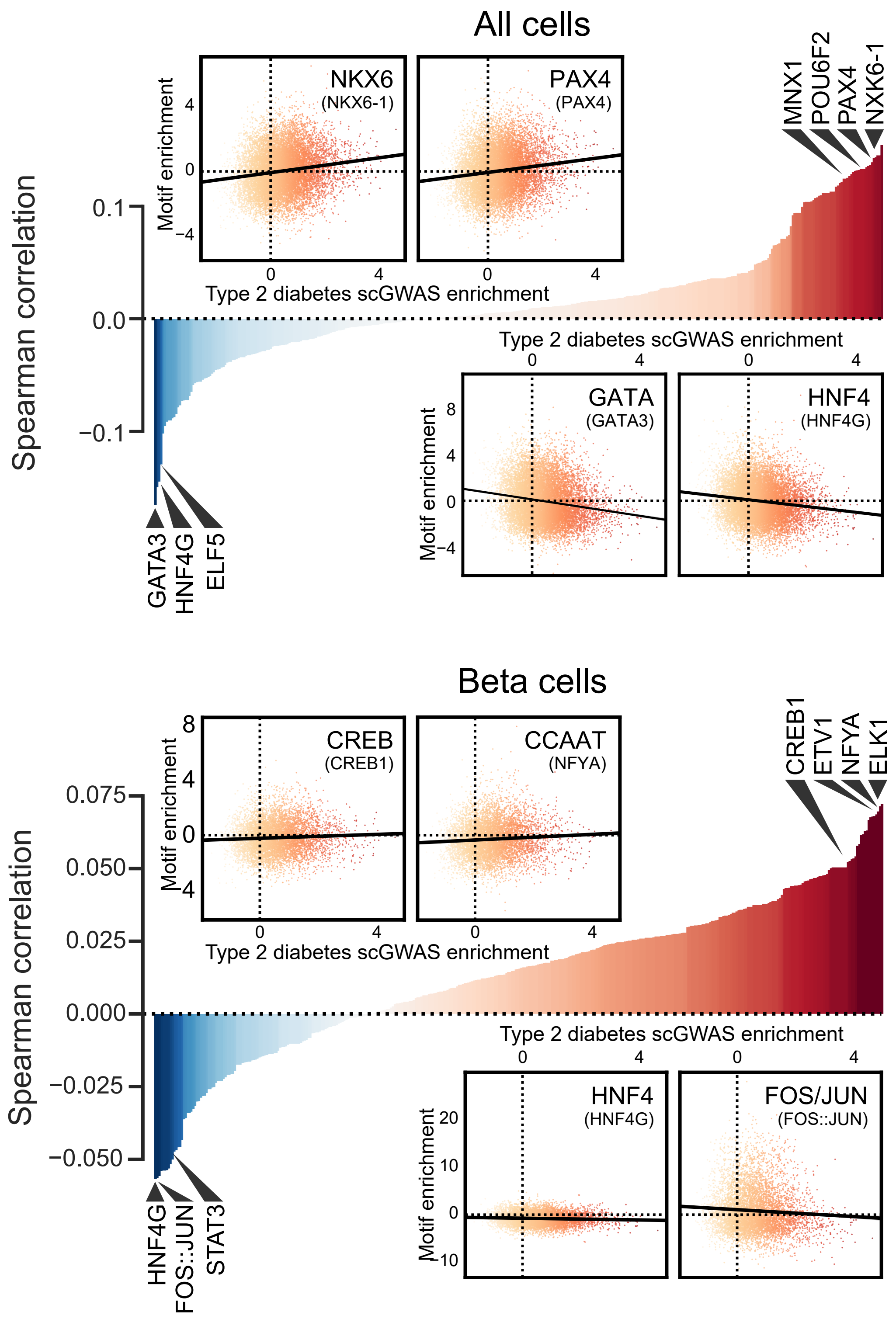


**Supplementary Figure 9. Single cell enrichment for type 2 diabetes.** Spearman correlation (y-axis) between single cell T2D enrichments and motif enrichment z-scores using datapoints for all cells (top) and within beta cells only (bottom). Within all cells, motifs subfamilies with the highest correlations include NKX6 and PAX4 (positive correlations) and GATA and HNF4 (negative correlations). WIthin beta cells, motif subfamilies with the highest correlations include CREB and CCAAT (positive correlations) and FOS/JUN and HNF4 (negative correlations). Highlighted T2D-motif correlations in either direction are labeled and raw data points comparing T2D single cell GWAS (scGWAS) enrichment to motif enrichment z-scores are shown on scatterplots (insets). Motif subfamily names are displayed in larger text on the scatterplot with the source motif in parentheses in smaller text below.

### Figure S10

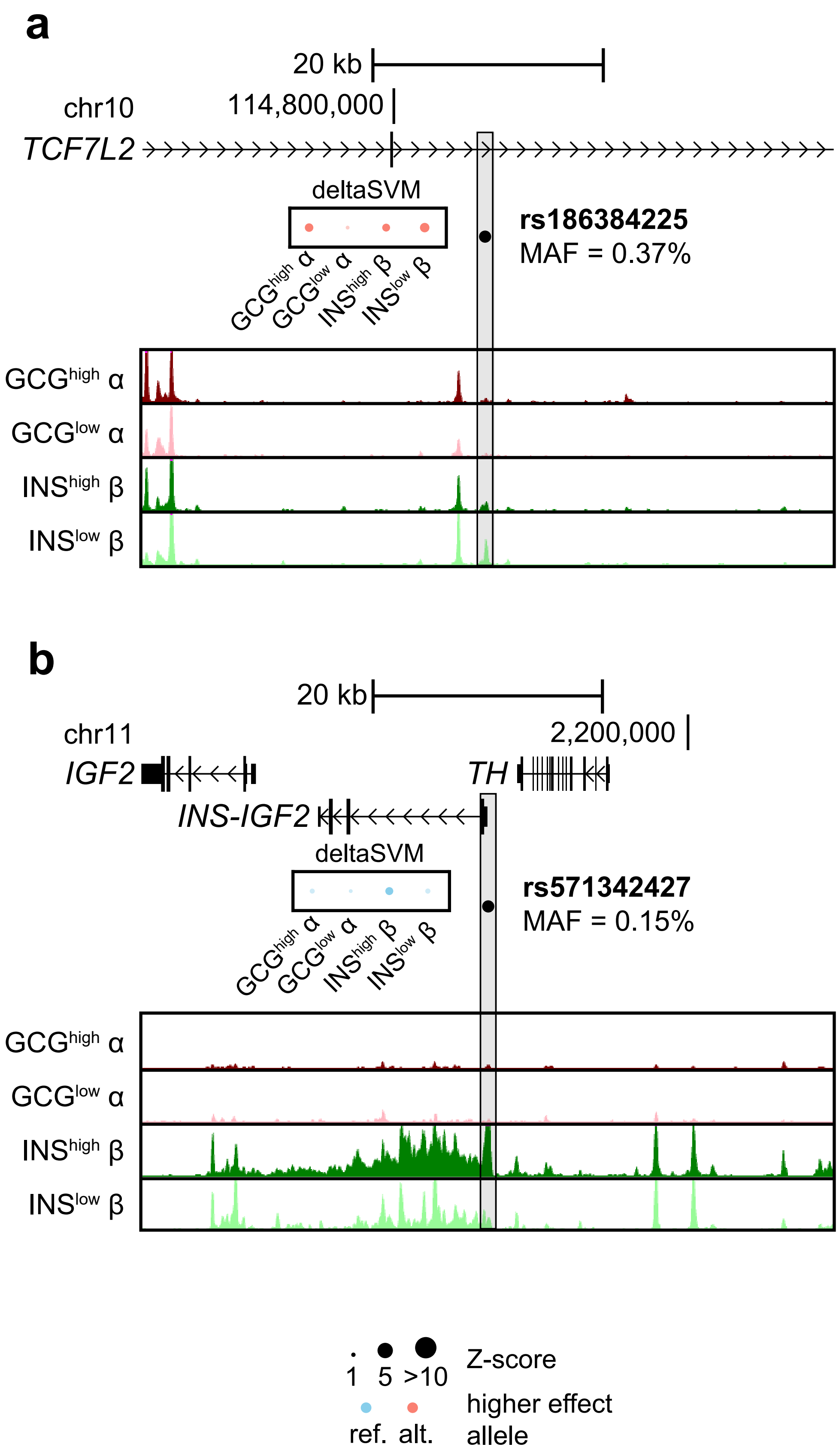


**Supplementary Figure 10. Rare genetic variants with predicted effects on chromatin accessibility.** (a) The rare variant rs186384225 (MAF=0.37% in DIAMANTE), located within an intron of *TCF7L2*, has significant deltaSVM effects with higher activity for the reference allele in $GCG^{high}$ α and $INS^{high}$ and $INS^{low}$ β cell states. (b) The rare variant rs571342427 (MAF=0.15%), located near the promoter of *INS-IGF2*, has significant deltaSVM effects for the alternative allele in $INS^{high}$ β cells.

### Figure S11

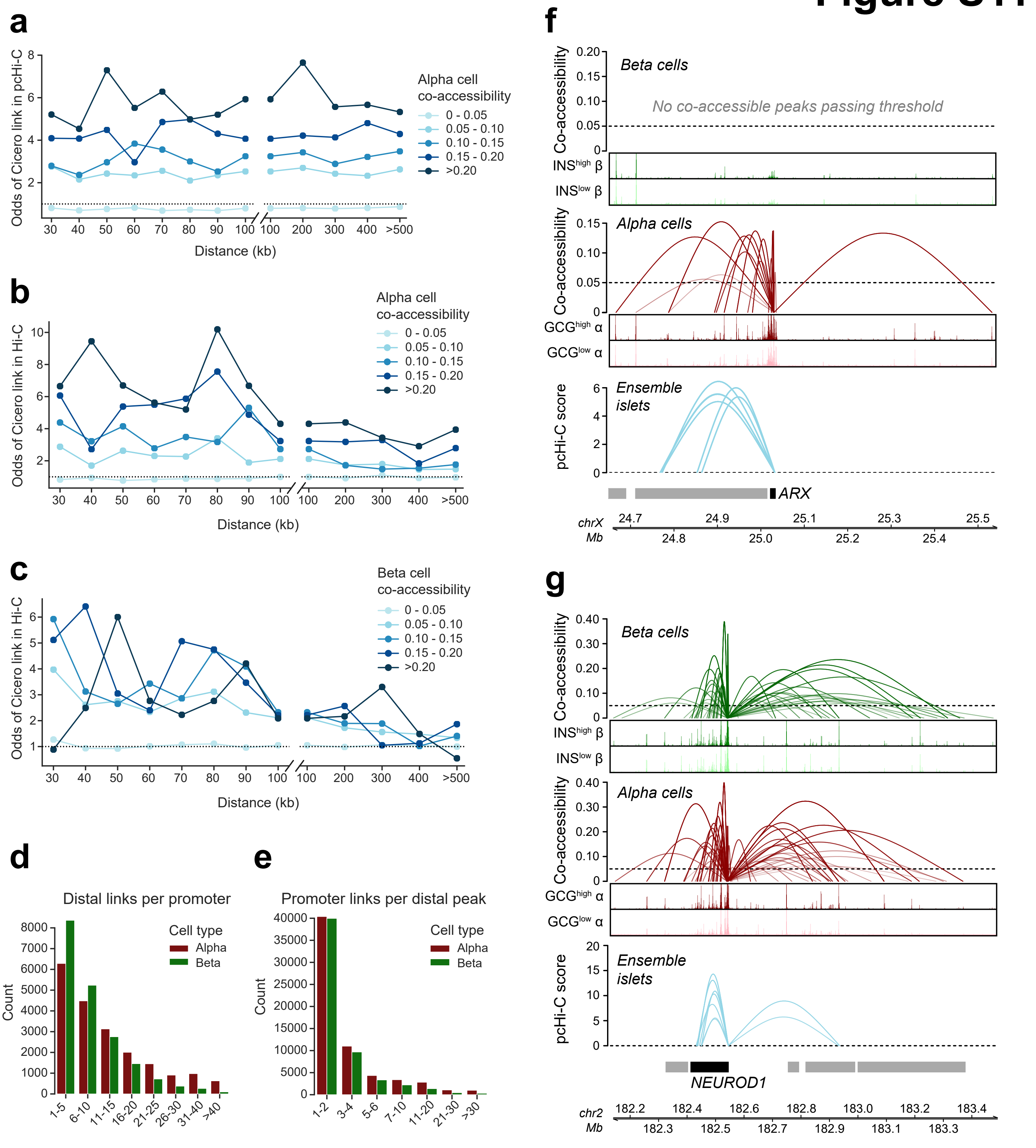

a
Odds of Cicero link in pcHi-C
Distance (kb)
Alpha cell co-accessibility
0 - 0.05
0.05 - 0.10
0.10 - 0.15
0.15 - 0.20
>0.20
b
Odds of Cicero link in Hi-C
Distance (kb)
Alpha cell co-accessibility
c
Odds of Cicero link in Hi-C
Distance (kb)
Beta cell co-accessibility
d
Distal links per promoter
Count
Cell type
Alpha
Beta
e
Promoter links per distal peak
Count
Cell type
Alpha
Beta
f
Co-accessibility
Beta cells
No co-accessible peaks passing threshold
INShigh β
INSlow β
Alpha cells
GCGhigh α
GCGlow α
pcHi-C score
Ensemble islets
ARX
chrX Mb
g
Co-accessibility
Beta cells
INShigh β
INSlow β
Alpha cells
GCGhigh α
GCGlow α
pcHi-C score
Ensemble islets
NEUROD1
chr2 Mb

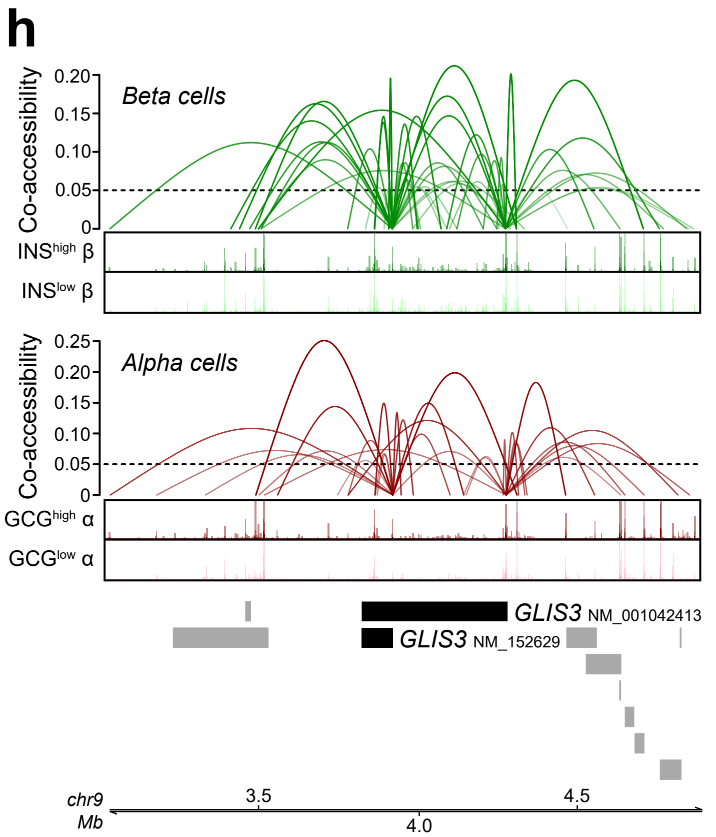


**Supplementary Figure 11. Single cell co-accessibility in alpha and beta cells.** (a) Distance-matched odds that alpha cell co-accessibility links overlap pcHi-C chromatin loops at different co-accessibility threshold bins in 0.05 intervals demonstrate that co-accessible links are enriched for chromatin interactions. (b) Same analysis as in A but with alpha cell co-accessibility and Hi-C loops. (c) Same analysis as A but with beta cell co-accessibility and Hi-C loops. (d) Number of distal sites linked to each promoter peak for alpha (red) and beta (green) cells. (e) Number of promoters linked to each distal site for alpha (red) and beta (green) cells. (f) An example of alpha cell-specific co-accessibility where beta (top) and alpha cell co-accessibility (middle) and high-confidence pcHi-C loops from ensemble islets (bottom) are anchored on the *ARX* promoter. (g) An example of shared co-accessibility where beta (top) and alpha cell co-accessibility (middle) and high-confidence pcHi-C loops from ensemble islets (bottom) are anchored on the *NEUROD1* promoter. (h) An example at the *GLIS3* gene which has at least two known transcript variants (Refseq transcript IDs: NM_001042413 and NM_152629) with accessible promoters and is co-accessible with distal sites with high hormone and low hormone dependent activity.

### Figure S12

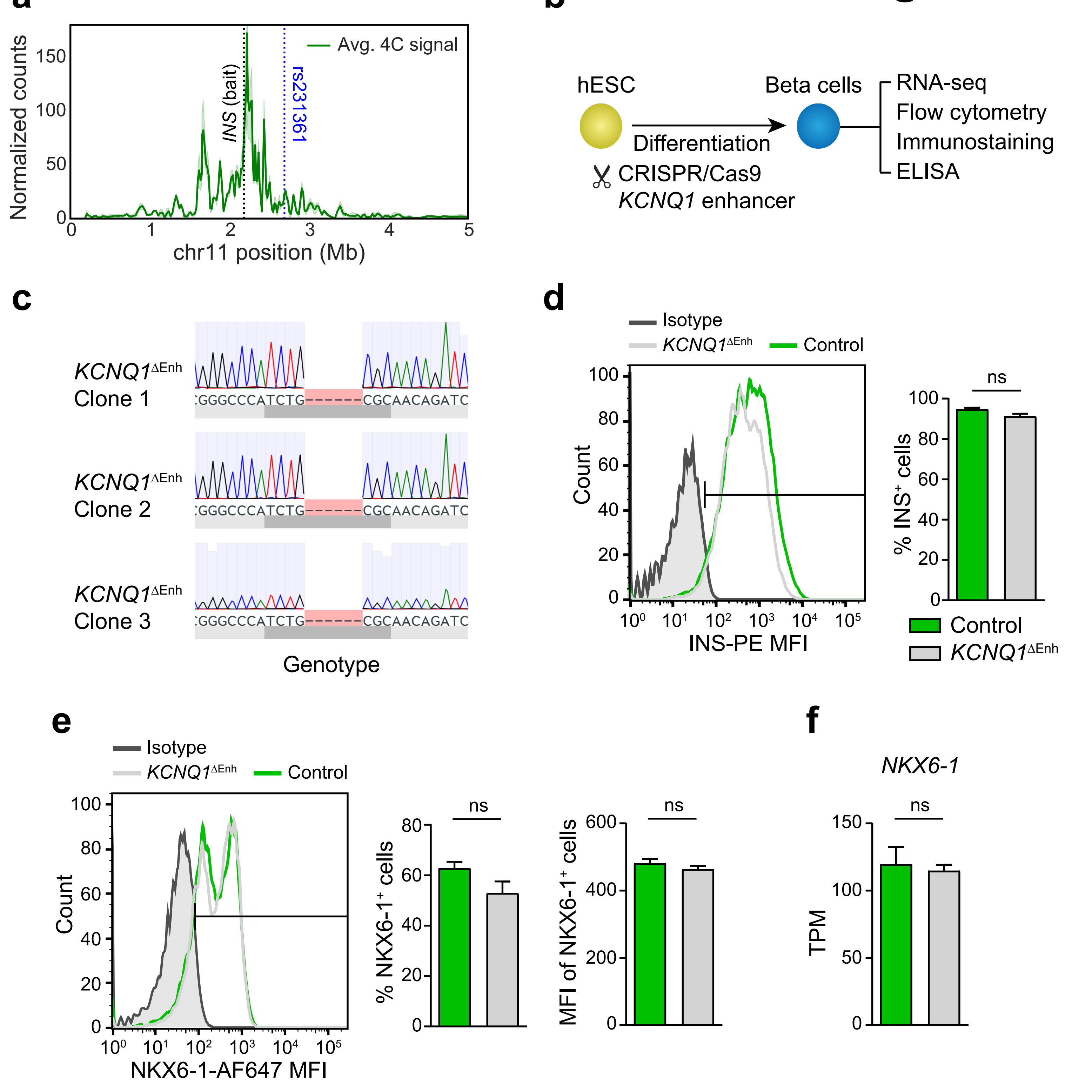


**Supplementary Figure 12. *KCNQ1* enhancer deletion in hESCs and effects on beta cell differentiation.** (a) Normalized 4C read counts (thousands) averaged from 3 replicates of EndoC beta cells (shaded line shows minimum and maximum from 3 replicates) using the *INS* promoter (black dotted line) as the bait region and highlighting rs231361. (b) Schematic of the workflow. (c) Sanger sequencing showing *KCNQ1* enhancer deletion in three independent hESC clones. (d) Histogram showing percentage of $INS^{+}$ cells by flow cytometry with a gating bar (left) and quantification (right) in beta cell stage cultures from $KCNQ1^{\Delta Enh}$ (n=9; 3 clones each differentiated three times) and control (n=6; 2 clones each differentiated three times) cells. (e) Histogram showing percentage of NKX6-$1^{+}$ cells with a gating bar (left) and quantification (middle) and median NKX6-1 fluorescence intensity (MFI, right) in beta cell stage cultures from $KCNQ1^{\Delta Enh}$ (n=9; 3 clones each differentiated three times) and control (n=6; 2 clones each differentiated three times) cells. (f) *NKX6-1* mRNA expression (in transcripts per million, TPM) in beta cell stage cultures from $KCNQ1^{\Delta Enh}$ (n=6; 3 clones each differentiated two times) and control (n=2; 1 clone differentiated two times) hESC clones. ns, not significant.

### Figure S13

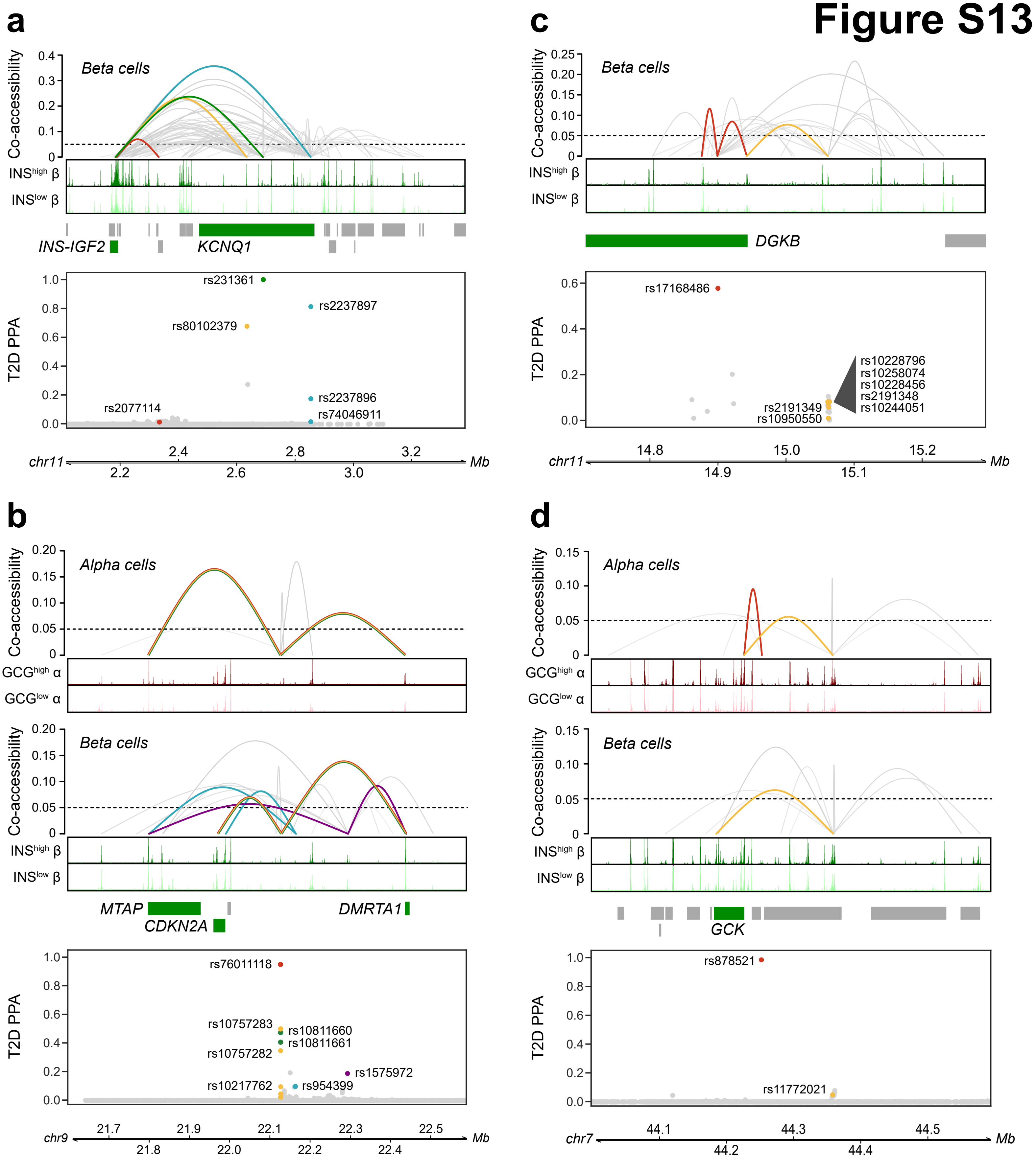


**Supplementary Figure 13. T2D risk variants in sites co-accessible with putative target gene promoters.** Multiple fine mapped T2D signals, including (a) four at the *KCNQ1* locus, (b) five at the *CKDN2A* locus, (c) two at the *DGKB* locus, and (d) two at the *GCK* locus overlap highly distal regulatory sites, are linked to promoters for the same genes through co-accessibility in alpha (top panels) and beta (middle panels) cells, and contain variants with high probability (PPA, bottom panels). Variants with PPA>.01 that overlap a distal linked site are labeled and colored separately for each independent signal, and their corresponding co-accessibility links to gene promoters are colored accordingly.

Figure S14

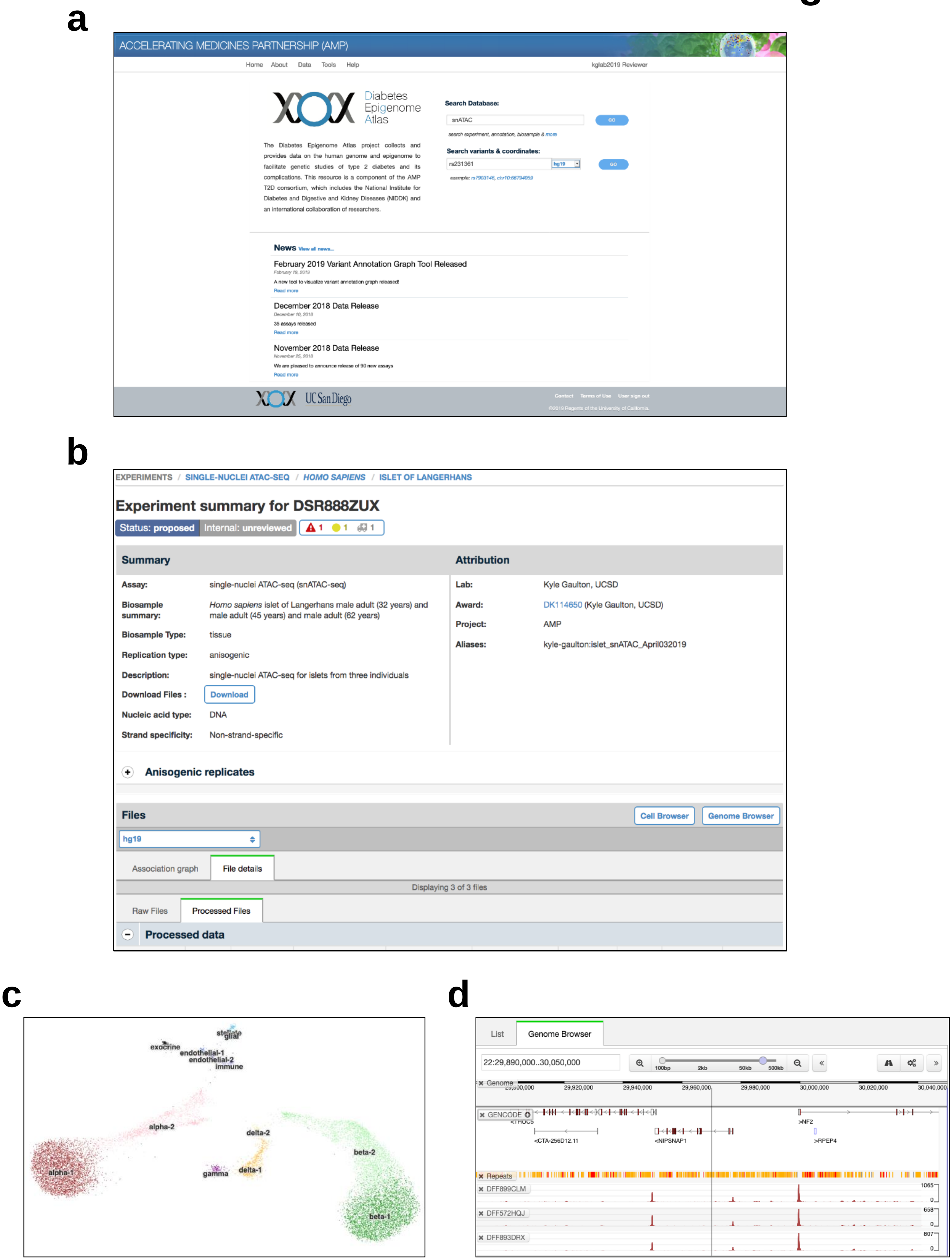
a
ACCELERATING MEDICINES PARTNERSHIP (AMP)
Home About Data Tools Help
kglab2019 Reviewer
Diabetes Epigenome Atlas
Search Database:
snATAC
GO
search experiment, annotation, biosample & more
Search variants & coordinates:
rs231361
hg19
GO
The Diabetes Epigenome Atlas project collects and provides data on the human genome and epigenome to facilitate genetic studies of type 2 diabetes and its complications. This resource is a component of the AMP T2D consortium, which includes the National Institute for Diabetes and Digestive and Kidney Diseases (NIDDK) and an international collaboration of researchers.
News View all news...
February 2019 Variant Annotation Graph Tool Released
A new tool to visualize variant annotation graph released!
Read more
December 2018 Data Release
35 assays released
Read more
November 2018 Data Release
We are pleased to announce release of 90 new assays
Read more
UC San Diego
Contact Terms of Use User sign out
b
EXPERIMENTS / SINGLE-NUCLEI ATAC-SEQ / HOMO SAPIENS / ISLET OF LANGERHANS
Experiment summary for DSR888ZUX
Status: proposed
Internal: unreviewed
Summary
Assay: single-nuclei ATAC-seq (snATAC-seq)
Biosample summary: Homo sapiens islet of Langerhans male adult (32 years) and male adult (45 years) and male adult (62 years)
Biosample Type: tissue
Replication type: anisogenic
Description: single-nuclei ATAC-seq for islets from three individuals
Download Files : Download
Nucleic acid type: DNA
Strand specificity: Non-strand-specific
Attribution
Lab: Kyle Gaulton, UCSD
Award: DK114650 (Kyle Gaulton, UCSD)
Project: AMP
Aliases: kyle-gaulton:islet_snATAC_April032019
Anisogenic replicates
Files
Cell Browser
Genome Browser
hg19
Association graph
File details
Displaying 3 of 3 files
Raw Files
Processed Files
Processed data
c
stellate
glial
exocrine
endothelial-1
endothelial-2
immune
alpha-2
delta-2
beta-2
alpha-1
gamma
delta-1
beta-1
d
List
Genome Browser
22:29,890,000..30,050,000
100bp 2kb 50kb 500kb
Genome
29,920,000 29,940,000 29,960,000 29,980,000 30,000,000 30,020,000 30,040,000
GENCODE
<THOC5
<CTA-256D12.11
<NIPSNAP1
>NF2
>RPEP4
Repeats
DFF899CLM
1065
0
DFF572HQJ
658
0
DFF893DRX
807
0

e

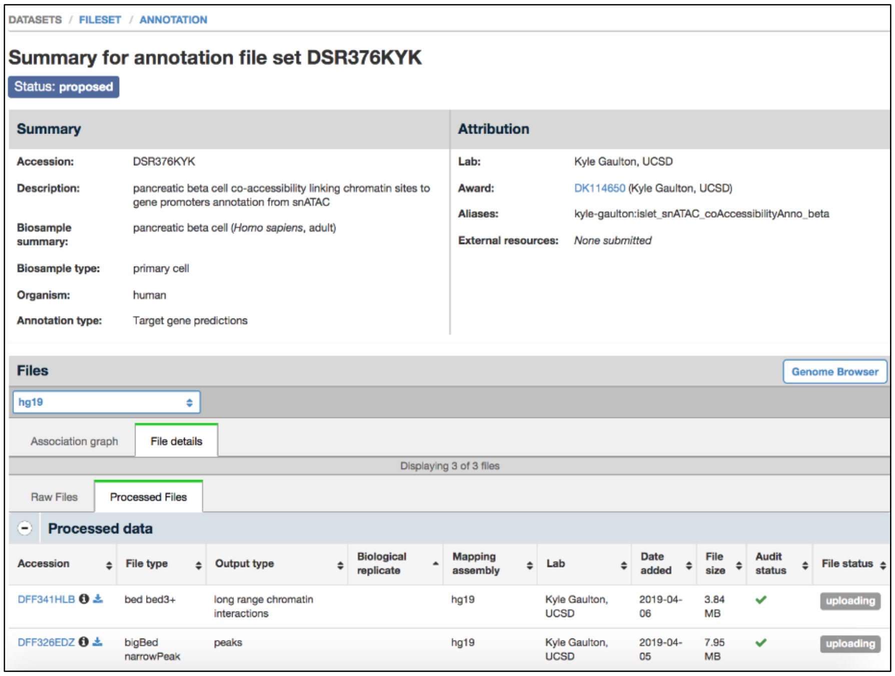


f

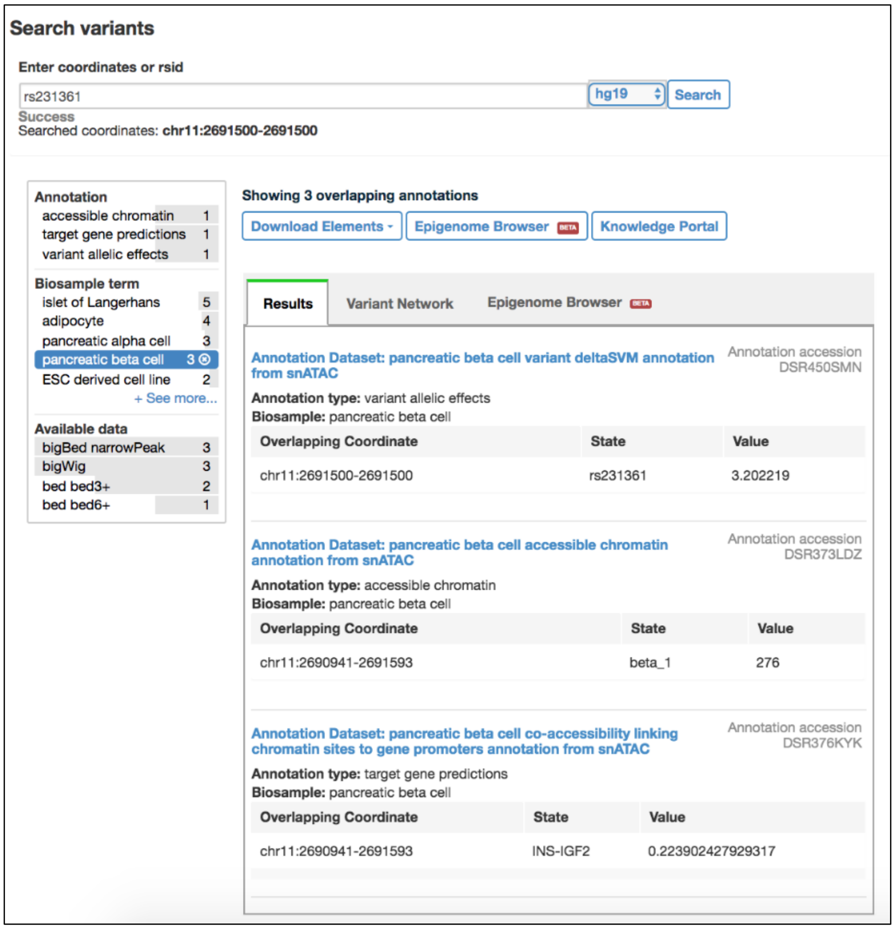


**Supplementary Figure 14. Web resource and database of islet snATAC-seq data.** (a) Interface of homepage where user can search for epigenome experimental and annotation files and query genetic variants for overlap with epigenomic annotations. (b) Interface of experiment page for islet snATAC-seq data from this study, which also provides links to view (c) cell projections, promoter accessibility patterns and TF motif enrichments in the UCSC cell browser and (d) signal tracks of accessible chromatin in the WashU Epigenome browser. (e) Interface of annotation page for single cell co-accessibility between distal sites and gene promoters in beta cells. (f) Interface of query for genetic variant overlap with annotations from islet snATAC-seq and other diabetes-relevant annotations. The variant rs231361 at the *KCNQ1* locus is shown in this example.
